# Telomeric 8-Oxoguanine Drives Rapid Premature Senescence In The Absence Of Telomere Shortening

**DOI:** 10.1101/2021.05.05.442662

**Authors:** Ryan P. Barnes, Mariarosaria de Rosa, Sanjana A. Thosar, Ariana C. Detwiler, Vera Roginskaya, Bennett Van Houten, Marcel P. Bruchez, Jacob Stewart-Ornstein, Patricia L. Opresko

## Abstract

Oxidative stress is a primary cause of cellular senescence and contributes to the pathogenesis of numerous human diseases. Oxidative damage to telomeric DNA is proposed to trigger premature senescence by accelerating telomere shortening. Here we tested this model directly using a precision tool to produce the common base lesion 8-oxo-guanine (8oxoG) exclusively at telomeres in human fibroblast and epithelial cells. A single induction of telomeric 8oxoG is sufficient to trigger multiple hallmarks of p53-dependent senescence. Telomeric 8oxoG activates ATM and ATR signaling, and enriches for markers of telomere dysfunction in replicating, but not quiescent cells. Acute 8oxoG production fails to shorten telomeres, but rather generates fragile sites and delayed mitotic DNA synthesis at telomeres, indicative of impaired replication. Based on our results we propose that oxidative stress promotes rapid senescence by producing oxidative base lesions which drive replication-dependent telomere fragility and dysfunction in the absence of shortening and shelterin loss.

## INTRODUCTION

Mammalian telomeres at chromosomal ends consist of long tandem 5’ TTAGGG 3’ arrays that are specifically bound by shelterin, a protein complex that remodels the end to suppress false activation of the DNA damage response (DDR) pathways and chromosome end-to-end fusions (de Lange, 2018). Progressive telomere shortening with cell division activates the DDR and triggers cellular” replicative senescence”, also called the Hayflick limit, characterized by irreversible cell cycle arrest and phenotypic changes (Bodnar et al., 1998; Hayflick and Moorhead, 1961). In this way, telomeres serve a potent tumor-suppressive function by limiting proliferative capacity (Greenberg et al., 1999). However, senescent cells accumulate with age and contribute to numerous ageing-related pathologies and diseases by compromising regenerative capacity and secreting inflammatory cytokines, chemokines and proteases that promote inflammation and alter the tissue microenvironment (Campisi et al., 2011; Hernandez-Segura et al., 2018). These changes in the microenvironment are also more permissive for tumor growth, and thus, paradoxically senescence can also promote tumorigenesis, metastasis, or immunosuppression (Kim et al., 2017; Krtolica et al., 2001; Kuilman et al., 2008; Ruhland et al., 2016). Furthermore, telomere dysfunction in pre-malignant cells with compromised DDR signaling, can cause chromosome end-to-end fusions and chromosomal instability which drives carcinogenesis (Artandi et al., 2000; Maciejowski et al., 2015). Thus, telomere function and integrity are critical for genome stability, cellular function and organism health.

A wealth of studies from human tissues, mice, and cell culture consistently show that chronic oxidative stress (OS) and chronic inflammation are associated with accelerated telomere shortening and dysfunction (Ahmed and Lingner, 2018; Barnes et al., 2019). Oxidative stress occurs when reactive oxygen species (ROS) arising from endogenous or exogenous agents, exceed antioxidant defenses, and can promote senescence and the pathogenesis of numerous degenerative diseases that occur with aging (Hegde et al., 2012; Malinin et al., 2011; Reuter et al., 2010). Guanine is the most susceptible to ROS of all the bases, and TTAGGG repeats are preferred sites for production of the minor, oxidative base lesion 8-oxoguanine (8oxoG) (Henle et al., 1999; Oikawa et al., 2001; Steenken, 1997). The links between OS and telomere shortening led to a model, proposed nearly 20 years ago, that oxidative modification to telomeric bases may contribute much greater to telomere loss and telomere-driven senescence, than the end-replication problem (von Zglinicki, 2002). ROS-induced damage to telomeres has also frequently been invoked to explain telomere dysfunction arising in low-proliferative tissues such as lung and heart, and arising independently of telomere length in mice with aging and chronic inflammation (Anderson et al., 2019; Birch et al., 2015; Jurk et al., 2014; Lewis-McDougall et al., 2019). A recent study showed that infiltrating neutrophils in liver trigger senescence in neighboring hepatocytes by ROS transmission, which generates telomere dysfunction in the absence of shortening (Lagnado et al., 2021). In these studies, dysfunctional telomeres are recognized by the localization of phosphorylated histone H2AX (γH2AX) and 53BP1 at telomeres, which are downstream effectors of DDR signaling kinases ATM and ATR (d’Adda di Fagagna et al., 2003; Sfeir and de Lange, 2012). These markers are referred to as telomere dysfunction induced foci (TIF) or telomere-associated DDR foci (TAF). While telomere deprotection upon genetic depletion of shelterin also activates the DDR (Sfeir and de Lange, 2012), evidence is lacking that OS-induced modification of telomeric DNA is extensive enough to completely displace shelterin and/or induce telomere deprotection. Therefore, the precise mechanism of ROS-induced DDR activation at telomeres, and whether oxidative modification of telomeric DNA can directly trigger senescence, remain unknown.

Delineating the biological impact of oxidative base lesions at telomeres has been challenging because oxidants used to modify DNA have pleiotropic effects on cell signaling, redox status, and transcription. To examine a direct causal role for oxidative base modifications in telomere loss, we developed a FAP-TRF1 chemoptogenetic tool that produces 8oxoG exclusively at telomeres with high spatial and temporal control via singlet oxygen, in the absence of damage elsewhere and redox changes (Fouquerel et al., 2019). 8oxoG is one of the most common base lesions and its physiological importance in mammalian cells is underscored by the evolution of three dedicated enzymes that specifically recognize this base in various contexts to enable repair or prevent mutations (Markkanen, 2017). Using the FAP-TRF1 tool to produce telomeric 8oxoG in human cancer cells, we found that acute lesion production has no obvious effects, but mimicking chronic OS with repeated telomeric 8oxoG formation over a month accelerates telomere shortening and losses. The accumulation of telomeric 8oxoG in cells lacking OGG1 enzyme which removes 8oxoG, increases hallmarks of telomere crisis including chromosomes fusions and chromatid bridges. This study, and work from others, provide evidence that accumulation of unrepaired 8oxoG can drive telomere instability in human cancer cells (Baquero et al., 2021; Fouquerel et al., 2019). However, a potential role for telomeric 8oxoG in cellular senescence and aging could not be delineated in genetically unstable cancer cell lines.

Here we demonstrate that in stark contrast to human cancer cells, acute production of the minor base lesion 8oxoG in telomeres is sufficient to rapidly impair growth of non-diseased human fibroblast and epithelial cell lines possessing wild-type DDR signaling mechanisms. Using our FAP-TRF1 tool we find that a single five minute treatment to produce telomeric 8oxoG induced numerous hallmarks of cellular senescence rapidly within 4 days of recovery (reviewed in (Hernandez-Segura et al., 2018)). These hallmarks include increased beta-galactosidase expression and mitochondrial oxygen consumption, reduced S-phase cells, and compromised nuclear Lamin integrity with concomitant increases in cytoplasmic chromatin or micronuclei. Remarkably, even though telomeres are less than 0.025% of the genome, the production of telomeric 8-oxoG was sufficient to rapidly activate ATM and ATR kinases and downstream effectors p53 and p21 within 30 minutes following guanine oxidation. Knock-out of p53 rescued the growth reduction, indicating p53 signaling enforces the 8oxoG-induced premature senescence. We provide evidence that the mechanism of premature senescence is by 8oxoG provoking replication stress-induced DDR activation and fragility at telomeres, rather than by accelerating telomere losses or shortening. Telomere fragility is widely reported as indicative of impaired telomere replication (Martinez et al., 2009; Sfeir et al., 2009). Our data reveal a novel mechanism of rapid telomere-driven senescence triggered by a common oxidative base lesion, that is distinctly different from “replicative senescence”, and has important implications for cellular aging linked to oxidative stress.

## RESULTS

### Acute Telomeric 8oxoG Initiates Rapid Premature Senescence in Non-Diseased Cells

We previously validated the FAP-TRF1 system for specifically inducing 8oxoG at telomeres in cells, when treating with MG2I dye and 660nm red light together (Fouquerel et al., 2019). To understand how cells derived from non-diseased tissue respond to telomeric 8oxoG, we generated clones that homogenously express FAP-mCerulean-TRF1 (termed FAP-TRF1) at telomeres in the human fibroblast BJ-hTERT and epithelial RPE-hTERT cell lines (hereafter referred to as BJ and RPE FAP-TRF1) (Figures 1A-B). These cell lines were transduced with telomerase at an early passage making them amenable for cloning, but importantly, they exhibit normal karyotypes and DNA damage response pathways. To confirm 8oxoG induction, we transfected RPE FAP-TRF1 cells with YFP-XRCC1 and observed an increase in both XRCC1 co-localization events and signal intensity at telomeres after 100 nM dye and 10 minute light treatment (Figures S1A-C). This treatment did not displace FAP-TRF1 from the telomeres, as evidenced by no loss of telomeric mCerulean foci (Figure S1D). The increase in XRCC1 at telomeres was attenuated in OGG1 knock-out or sodium azide treated cells, verifying XRCC1 recruitment is specific to 8oxoG generated by singlet oxygen (Figures S1B-C, S1E-F). We investigated how induction of the minor base lesion 8oxoG at telomeres impacts the growth of non-diseased cells. In contrast to untreated cells or treatment with dye or light alone, BJ and RPE FAP-TRF1 cells treated with dye and light together for five minutes showed a highly significant reduction in cell growth just 4 days after treatment (Figure 1C-D, and S1G for additional RPE FAP-TRF1 clone). Importantly, the extent of growth reduction was dependent on the duration of light exposure, showing the cellular response is proportional to the amount of telomeric damage (Figure S1H). When we repeated this experiment in the parental hTERT BJ and RPE cells not expressing FAP-TRF1, and in FAP-TRF1 expressing HeLa and U2OS cancer cells, we observed no change in growth (Figures S1I-K). These data show the growth reduction observed in BJ and RPE FAP-TRF1 cells with dye and light treatment requires FAP-TRF1 expression, and confirms cancer cells are unaffected by an acute treatment (Fouquerel et al., 2019).

**Figure 1.**
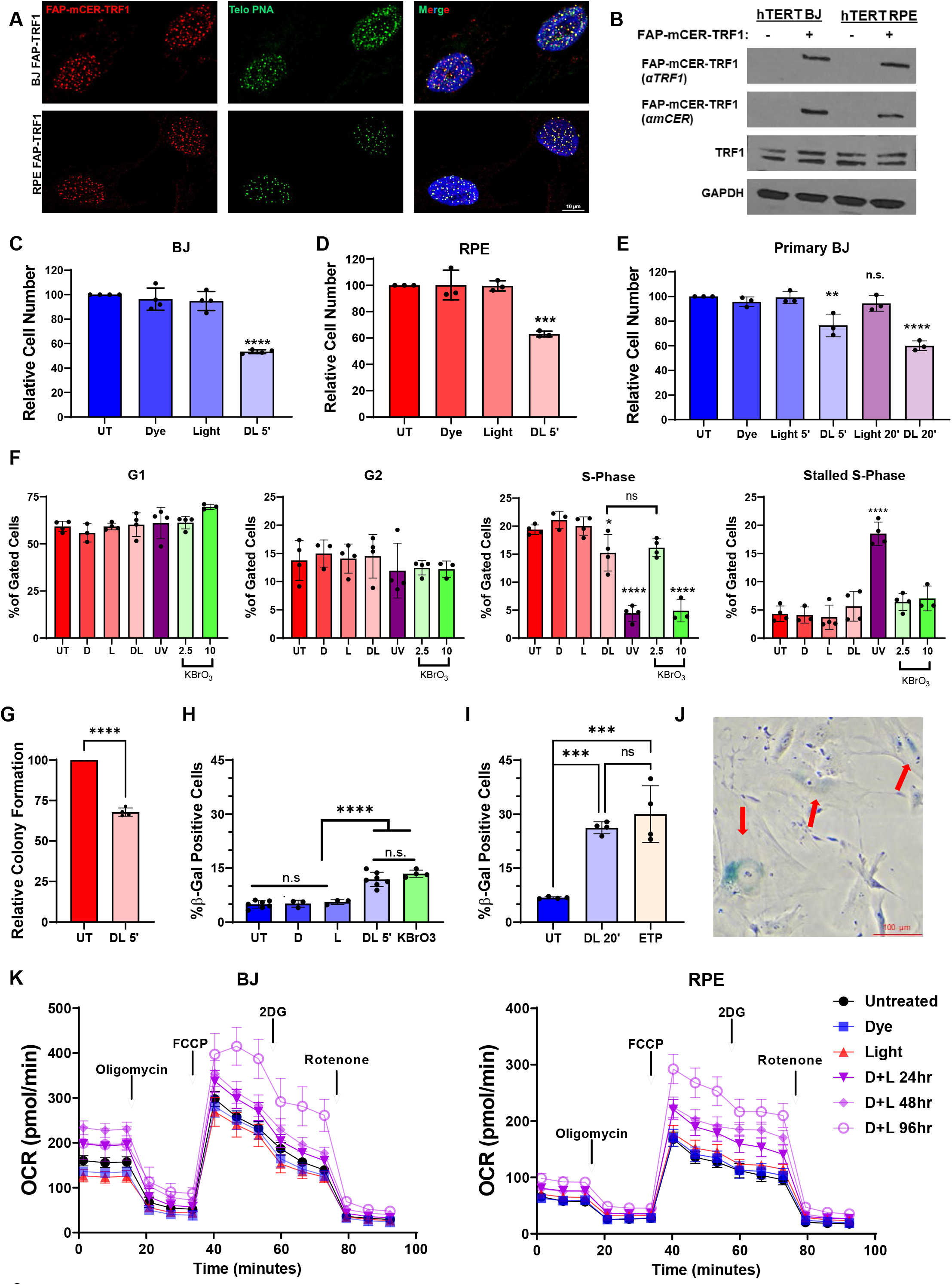
Acute Telomeric 8oxoG Initiates Rapid Premature Senescence in Non-Diseased Cells. (A) Representative images of FAP-mCER-TRF1 protein colocalization with telomeres in BJ (top panel) and RPE (bottom) clones expressing FAP-TRF1 visualized by anti-mCER staining (red) and with telo-FISH (green). (B) Immunoblot for TRF1 in whole cell extracts from hTERT BJ and RPE cells with and without FAP-mCER-TRF1 expression. TRF1 antibody detects both exogenous and endogenous TRF1 while mCER antibody detects exogenous only. (C-E) Cell counts of BJ (C), RPE (D), or primary BJ (E) FAP-TRF1 cells obtained 4 days after recovery from dye and light alone, or in combination (DL), relative to untreated (UT) cells. (F) Percent of RPE FAP-TRF1 cells at various cell cycle phases 24 hours after no treatment or exposure to dye, light, dye + light, 20 J/m^2^ UVC, or 1 hour treatment with 2.5 or 10 mM KBrO_3_ determine by flow cytometry. Cells were pulsed with EdU 1 hour before harvest. (G) Percent colony formation of RPE FAP-TRF1 cells obtained 7-10 days after treatment. (H-I) Percent β-galactosidase positive BJ FAP-TRF1 cells obtained 4 days after recovery from the indicated treatments. 2.5 mM KBrO_3_ and 50 μm etoposide (ETP) treatment was for 1 hour. (J) Representative image of 5 min dye and light treated BJ FAP-TRF1 β-galactosidase positive cells. Arrows mark positive cells (turquoise). (K) BJ and RPE FAP-TRF1 cells were treated for 5 min with dye, light or dye + light. Mitochondrial respiration was examined by OCR as measured with a Seahorse Extracellular Flux Analyzer 24, 48 and 96 h after dye + light. For panels C-I, data are means ± SD from at least three independent experiments and One-way ANOVA was performed, except for G which was t-test. For panel K, data are from 2 independent experiments with 7-8 technical replicates, and data are the mean ± 95% C.I. *p<0.05, **p<0.01, ***p<0.001, ****p<0.0001.

To determine whether the growth reduction depends on hTERT expression, we infected primary BJ cells with FAP-TRF1 lentivirus and grew them under selection without clonal expansion, resulting in stable but variable expression (Figure S1L). When these cells were analyzed by the same growth assay, we observed a consistent and dose dependent reduction in growth with 5 and 20 minutes of light and dye exposure (Figure 1E). Thus, induction of telomeric 8oxoG elicits growth reduction in non-diseased cells, regardless of telomerase status. To compare targeted formation of 8oxoG at telomeres to induction in the bulk genome, we used potassium bromate (KBrO_3_), which primarily produces 8oxoG, but also damages other cellular compartments (Parlanti et al., 2012). Interestingly, 2.5 mM KBrO_3_ treatment for one hour reduced BJ and RPE cell growth to levels comparable with five minutes dye and light treatment (Figures S1M and S1N). Remarkably, these data demonstrate that even though telomeres are a tiny fraction of the genome (<0.025%), 8oxoG confined to the telomeres reduces growth of non-diseased cells to levels comparable to 8oxoG produced in the bulk genome.

Next, we asked whether the reduction in growth triggered by telomeric 8oxoG formation was due to cellular senescence, which is characterized by persistent growth arrest in addition to other cellular and molecular phenotypes that vary depending on the cell type and the mechanism of senescence induction, i.e. replicative exhaustion or DNA damage induced premature senescence (Hernandez-Segura et al., 2018). We analyzed the cell cycle by flow cytometry 24 hours after dye and light treatment, since impaired cell growth was already apparent at this early-timepoint (Figure S1O). Following 5 minutes of light and dye treatment, RPE FAP-TRF1 cells showed a significant reduction in EdU positive S-phase cells, and a slight increase in stalled S-phase cells (EdU negative) (Figures 1F and S1P). These changes are comparable to 2.5 mM KBrO_3_, consistent with the growth data, while 10 mM KBrO_3_ and 20 J/m^2^ UVC exposures showed dramatic reductions in S-phase cells.

Senescence-associated (SA) growth arrest is most commonly assayed by colony formation and beta-galactosidase (β-gal) detection. For RPE FAP-TRF1 cells, we assessed colony formation to determine the fraction of cells that retain proliferative capacity following treatment. Cells were treated, and then plated at very low density to allow individual colonies to form. After 7-10 days of recovery, telomeric 8oxoG significantly reduced colony formation efficiency (Figure 1G). We stained BJ FAP-TRF1 cells for the expression of SA-β-gal, a well-established marker of senescent fibroblasts, 4 days after inducing telomere damage. Consistent with the growth assay, we observed a significant increase in β-gal positive cells after treatment with 5 minutes of dye and light together, but not dye or light alone (Figures 1H, 1J and S1Q). The increase in β-gal staining after dye and light exposure was identical to 2.5 mM KBrO_3_, consistent with the similar reduction in cell number (Figure S1M). When telomeric damage was increased by treatment with dye and light for 20 minutes, we observed a dramatic increase in β-gal staining, similar to that achieved with the genotoxic control etoposide, and consistent with the greater growth inhibition (Figures 1I and S1H, S1M, and S1Q). These data demonstrate formation of telomeric 8oxoG inhibits the proliferation of individual cells.

Senescent cells remain metabolically active despite their non-proliferative state. To determine if growth arrested BJ and RPE FAP-TRF1 cells maintain metabolic activity, we analyzed their mitochondria for oxygen consumption rate (OCR) and extra-cellular acidification rate (ECAR). Using the Seahorse technology 24, 48, and 96 hours after dye and light treatment, we found slight increases in the basal OCR, especially for the BJ FAP-TRF1 cells (Figure 1K). Significantly however, when the cells were treated with mitochondrial uncoupler FCCP, we observed a dramatic increase in the maximal respiration of cells treated with dye and light until the mitochondria were inhibited with rotenone (Figure 1K). No significant changes were observed for ECAR (data not shown). Our results are consistent with previous reports of elevated OCR in hydrogen peroxide or H-RAS induced senescence (Kim et al., 2018; Takebayashi et al., 2015). In summary, our data show non-diseased cells undergo rapid, premature senescence following telomeric 8oxoG formation.

### Telomeric 8oxoG Production Increases Cytoplasmic DNA

A shared hallmark of senescence and cancer is an increase in micronuclei, also termed cytoplasmic chromatin fragments (CCF), that can arise by different mechanisms (Hernandez-Segura et al., 2018; Krupina et al., 2021). 4 days after 5 minutes of dye and light exposure, both BJ and RPE FAP-TRF1 cells showed significant increases in micronuclei (Figure 2A-C). The majority of micronuclei stained positive for γH2AX and telomere DNA (>75%), but negative for 53BP1 (Figure 2A), consistent with previous reports for CCFs generated during replicative or premature senescence (Ivanov et al., 2013). Micronuclei of senescent cells are sensed by the cytoplasmic DNA sensor, cGAS, and this promotes the senescence associated secretory phenotype (SASP). Although 4 days is too soon for canonical SASP to develop, we observed a significant increase in cGAS positive micronuclei after dye and light treatment, while no change was observed for γH2AX positive micronuclei (Figure 2D). However, the percent of micronucleated cells positive for γH2AX increased with induction of telomeric 8oxoG (Figure 2E), demonstrating the increase in total and cGAS positive micronuclei is due to DNA damage.

**Figure 2.**
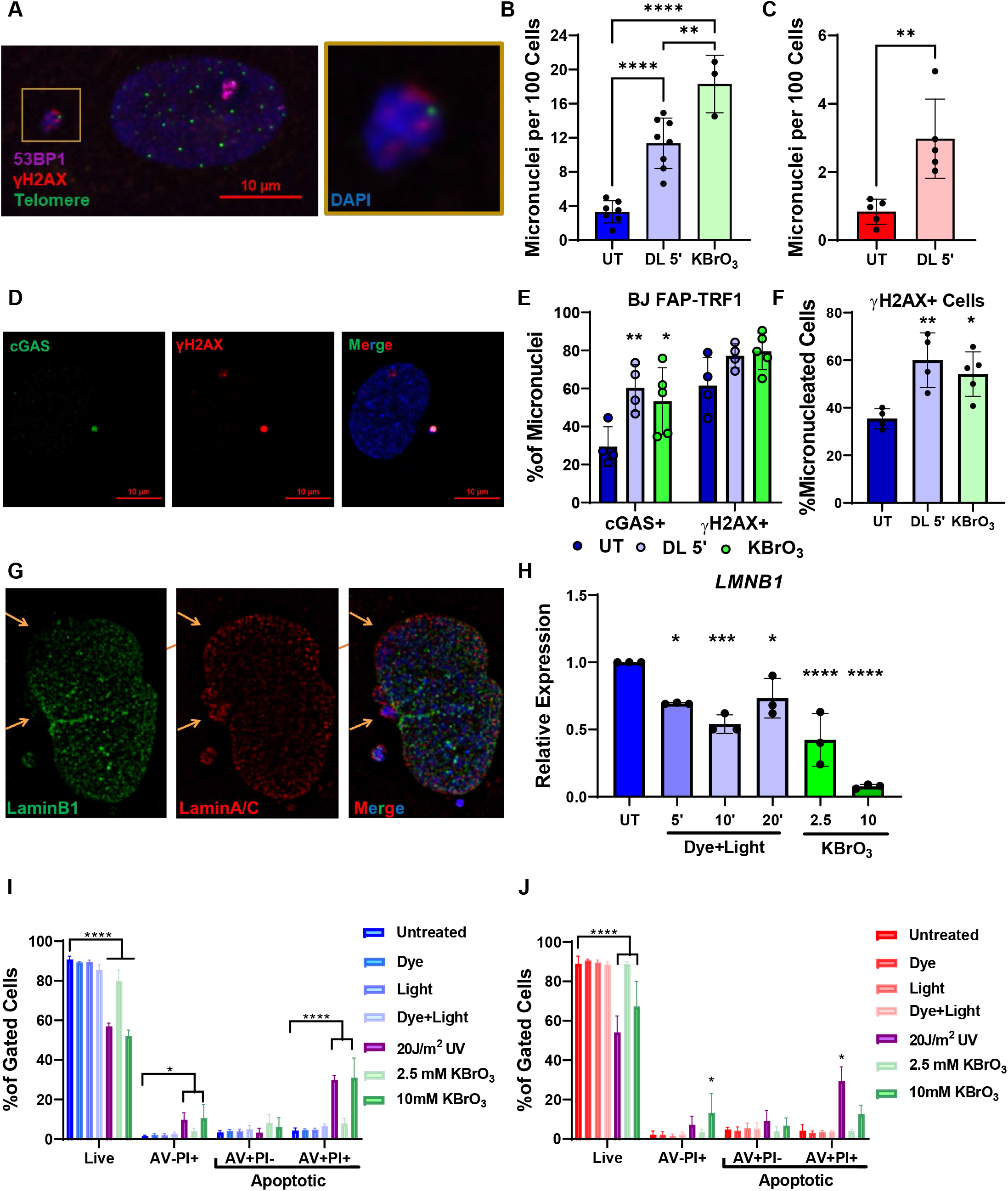
Telomeric 8oxoG Production Increases Cytoplasmic DNA. (A) Representative IF-teloFISH of micronuclei in BJ FAP-TRF1 cells 4 days after recovery from 5 min dye + light. Gold box indicates a zoomed in image of the micronuclei. (B-C) Quantification of number of micronuclei per 100 nuclei for BJ (B) or RPE (C) FAP-TRF1 cells 4 days after recover from 5 min dye + light, or for BJ after 1 h 2.5 KBrO_3_. (D) Representative IF of cGAS and γH2AX staining 4 days after 5 min dye and light treatment. (E) Quantification of the percent of micronuclei that are cGAS or γH2AX positive 4 days after 5 min dye + light or 1 hour 2.5 mM KBrO_3_ treatment. (F) Quantification of the percent of cells with micronuclei that are positive for γH2AX in the main nucleus. (G) Representative IF of BJ FAP-TRF1 cells for LaminB1 and LaminA/C obtained 4 days after 5 min dye + light treatment. Arrows point to absent LaminB1 staining where LaminA/C is intact. See Figure S2E. (H) Quantification of *LMNB1* mRNA from BJ FAP-TRF1 cells 4 days after 5 min light alone, 5, 10 or 20 min dye + light, or 1 h 2.5 or 10 mM KBrO_3_, relative to untreated. (I-J) Percent of BJ (H) or RPE (I) FAP-TRF1 cells positive for annexin V (AV), propidium iodide (PI), or both 4 days after indicated treatments. KBrO_3_ exposure was for 1 hour. For all graphs, data are means ± SD from at least three independent experiments, and statistical results indicated as *p<0.05, **p<0.01, ***p<0.001, ****p<0.0001. Panel C was analyzed by unpaired T-test, and panels B, E, F, H-J by One or Two-way ANOVA.

Micronuclei can arise from blebbing from the main nucleus due to altered chromatin and lamina structure, or due to aberrant mitosis. Immuno-fluorescence staining revealed that approximately 9% of treated cells displayed defects in Lamin B1 protein, where Lamin A/C was present (Figures 2G arrows and S2A). Consistent with this, we found reduced Lamin B1 mRNA and protein levels (Figures 2H and S2B). In addition, BJ and RPE FAP-TRF1 cells both showed significant increases in nuclei size 4 days after dye and light treatment (Figure S2C). We also scored for chromatin bridges, indicative of mitotic defects that can lead to micronucleus formation. However, these events were exceedingly rare occurring on average in 0.5% and 0.75% of treated BJ and RPE FAP-TRF1 cells, respectively. Together, these data indicate telomere 8oxoG induced micronuclei likely arise by a mechanism consistent with nuclear blebbing due to Lamin changes, and not chromatin bridge breakage.

We further investigated whether 8oxoG-induced micronuclei arise from DNA breaks in two ways. DNA damage induced apoptosis can directly produce breaks or result from activation of endonucleases. We tested for apoptosis by Annexin V (AV) and propidium iodine staining and flow cytometry (Figure S2D). 4 days after treatment with 20 J/m^2^ UV or 10 mM KBrO_3_, BJ and RPE FAP-TRF1 cells displayed late apoptotic (AV+PI+) and dead cells (AV-/PI+) cells, serving as positive controls (Figure 2I-J). Dye and 5 minute light treatment however, did not increase cell death or apoptosis compared to controls for either cell line. Interestingly, 2.5 mM KBrO_3_ treated cells again behaved similarly to dye and light treatment, suggesting a comparable cellular response from 8oxoG production in just the telomeres, to lesion production throughout the entire genome. We also directly tested for DNA breaks by performing electrophoresis of cells in agarose plugs (Figure S2E). Immediately following, or 24 hours after, treatment with dye and light for 5 minutes, we observed no elevation of DNA breaks above background, whereas treatment with hydrogen peroxide (H_2_O_2_) markedly induced DNA breaks. These observations show induction of telomeric 8oxoG does not induce DNA breaks directly, consistent with our previous report (Fouquerel et al., 2019), nor subsequent apoptosis.

### p53 DNA Damage Signaling is Required for 8oxoG Induced Senescence

Signaling through the DNA damage response (DDR) can drive cell cycle arrest and growth inhibition leading to senescence if the damage is extensive or unresolved (Hernandez-Segura et al., 2018). We observed activation of the ATM/Chk2 pathway within minutes following dye and light treatment (Figure 3A). This result is striking because small base modifications are not canonically associated with ATM activation (Chou et al., 2015; Ciccia and Elledge, 2010). To test the role of DDR signaling in 8oxoG induced senescence, we treated cells with dye and light and then cultured them with the ATM inhibitor (ATMi) KU60019 (Golding et al., 2009). Compared to DMSO controls, the ATMi treated cells displayed a rescue of both the damage induced colony formation and β-gal phenotypes (Figures 3B and 3C), confirming the role of ATM in telomeric 8oxoG induced senescence.

**Figure 3.**
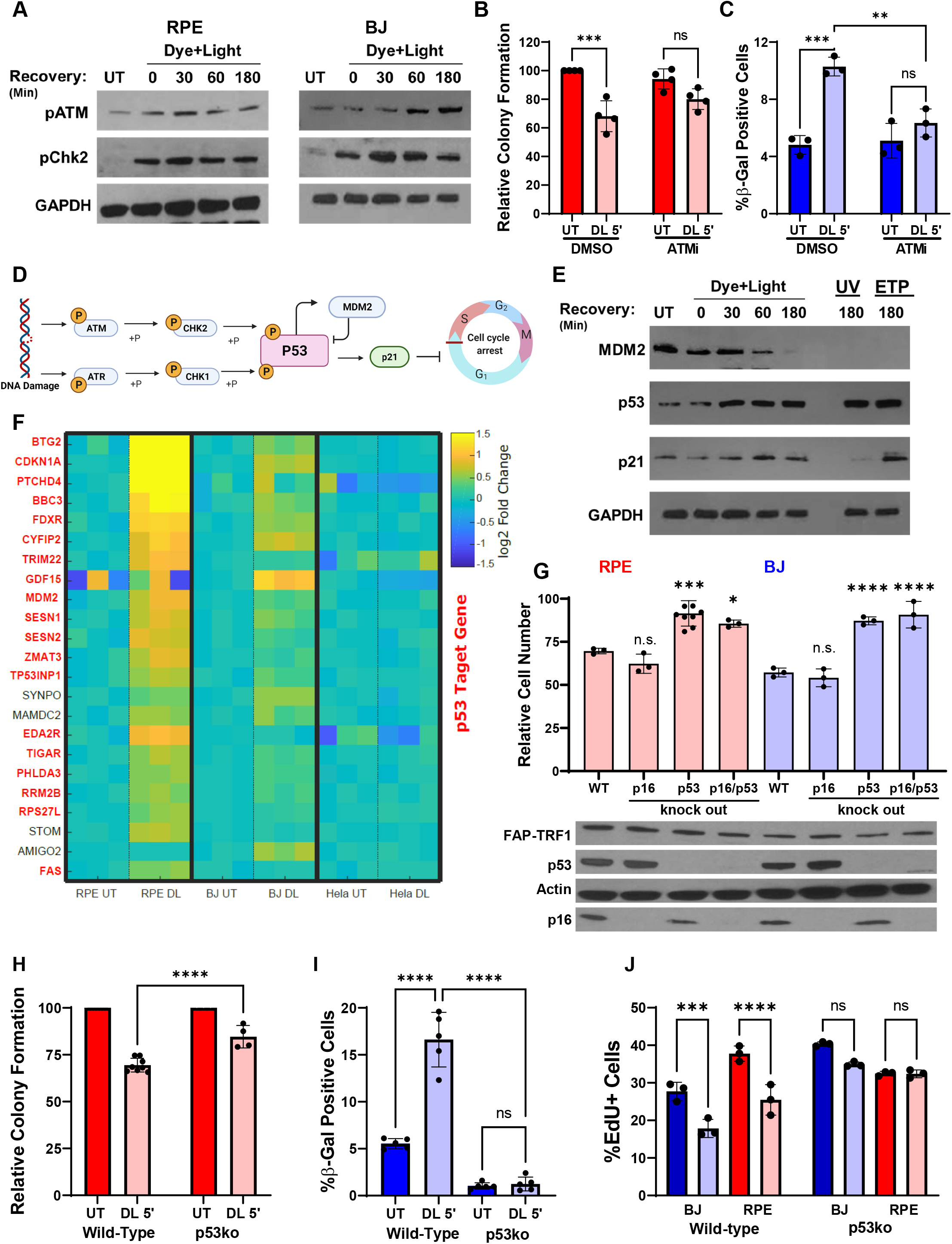
p53 DNA Damage Signaling is Required for 8oxoG Induced Senescence. (A) Immunoblots of phospho-ATM (pATM) and phospho-Chk2 (pChk2) from untreated BJ and RPE FAP-TRF1 cells, or cells treated with 5 min dye + light and recovered for the indicated times. (B) Colony formation of RPE FAP-TRF1 cells after recovery from indicated treatments. Cells were only cultured with ATMi or DMSO during recovery. (C) Percent β-gal positive BJ FAP-TRF1 cells obtained 4 days after recovery from indicated treatments. Cells were only cultured with ATMi or DMSO during recovery. (D) Schematic of canonical DNA damage induced p53 activation by ATM and ATR kinases. Created in BioRender. (E) Immunoblot of untreated BJ FAP-TRF1 cells, or cells treated with 5 min dye + light and recovered for the indicated times. UV = 20 J/m^2^ UVC. ETP = 1 hour 50 μM etoposide treatment. (F) Heat map of mRNA-Seq results from FAP-TRF1 expressing RPE, BJ, and HeLa cells 24 h after no treatment (NT) or 5 min dye + light (DL). Shown are significantly altered genes expressed in all three cell lines, p53 target genes are indicated in magenta. Each column is an independent replicate. (G) Cell counts of wild-type, p16ko, p53ko, or p16 + p53 double ko of BJ (blue) or RPE (red) FAP-TRF1 cells 4 days after recovery from 5 min dye + light treatment relative to untreated. Immunoblot below shows FAP-TRF1, p53, p16 expression with actin loading control. (H) Percent colony formation of wild-type or p53 ko RPE FAP-TRF1 cells obtained 4 days after recovery from 5 min dye + light treatment relative to untreated. (I) Percent β-gal positive wild-type or p53 ko BJ FAP-TRF1 cells obtained 4 days after recovery from no treatment or 5 min dye + light. (J) Percent EdU positive cells obtained from 1 h EdU pulse after 23 hours recovery from 5 min dye + light treatment of wild-type and p53ko BJ or RPE FAP-TRF1. Over 200 cells were scored per condition in each experiment. For panels B-C and G-J data are means ± SD from at least three independent experiments, and statistical analyses was by One-or Two-way-way ANOVA. *p<0.05, **p<0.01, ***p<0.001, ****p<0.0001.

Next, we examined activation of tumor suppressor p53, which is downstream of ATM/Chk2 in the DDR and drives the transcription of numerous factors related to DNA repair, the cell cycle checkpoint and senescence enforcement (Figure 3D) (Kastenhuber and Lowe, 2017). Shortly after dye and light treatment, the p53 antagonist MDM2 is down-regulated, resulting in stabilization of p53 protein and induction of p21, a p53 target protein (Figure 3E). Activation of p53 and p21 prevents transcription of S-phase factors by reducing RB phosphorylation and inhibiting E2F transcription factors, which we indeed observed following induction of telomeric 8oxoG (Figure S3A). Consistent with ATM activating p53 in response to telomeric 8oxoG, we observed reduced p53 protein levels in cells treated with ATMi after dye and light exposure, compared to DMSO controls (Figure S3B and S3C).

To understand the global transcriptional response to telomeric 8oxoG, we performed whole transcriptome mRNA-sequencing on RPE, BJ, and HeLa FAP-TRF1 cells 24 hours after five minutes of dye and light treatment. Consistent with the above data and our previous report (Fouquerel et al., 2019), HeLa cells showed no significant changes after a single induction of 8oxoG at the telomere (Figure 3F and data not shown). However, both RPE and BJ FAP-TRF1 cells showed significant changes in gene expression, which did not correlate with proximity to the telomeres, demonstrating these changes were not an artifact of inducing DNA damage at the telomeres (Figures 3F and S3D-G). We interrogated pathway changes using the Hallmark gene set enrichment analysis, and found down-regulation of replication and cell cycle pathways consistent with senescence (Supplemental Table 1). In both cell lines, Hallmark indicated an upregulation of the p53 pathway, consistent with significant up-regulation of several of the same p53 targets in RPE and BJ FAP-TRF1 cells after treatment (Figure 3F) (Fischer, 2017).

In addition to p53, p16 is another classical driver of senescence via reduction of RB phosphorylation (Ohtani et al., 2004). Therefore, we asked if p53 and/or p16 are required to initiate senescence following 8oxoG damage, by using CRISPR-Cas9 to knock-out (ko) these genes alone and in combination in FAP-TRF1 clones. We repeated the 4-day growth assay and observed that p16ko did not rescue the growth reduction, while p53ko alone, or in combination with p16 did (Figure 3G). In agreement, RT-qPCR analysis showed no significant induction of p16 mRNA with dye and light treatment, while 10 mM KBrO_3_ did increase expression (Figure S3H). Compared to wild-type cells, p53ko cells displayed an attenuated reduction in cell growth as a function of light exposure time (Figure S3I). Consistent with a rescue of telomeric 8oxoG induced senescence, p53ko also suppressed the reduced colony formation and increased β-gal activity of treated RPE and BJ FAP-TRF1 cells respectively, and reduction in EdU incorporation for both cell lines (Figures 3H-J). Moreover, p53ko also rescued growth reduction in response to KBrO_3_ (Figure S3J).

p21 is a principal driver of p53 signaling, and we observed upregulation of both p21 protein by W blot and mRNA by the RNA-seq analysis (Figure 3E-F). Other factors such as NF-κB can upregulate p21 in response to DNA damage, so we compared wild-type and p53ko cells for induction of p21 by IF (Nicolae et al., 2018). Additionally, p21 upregulation in senescent cells should correlate with a lack of DNA replication, so these cells were scored for EdU incorporation. In both RPE and BJ FAP-TRF1 cells, 24 hours after treatment we found significant induction of p21 only in non-replicating wild-type cells, and this increase was lost in p53ko cells (Figures SK-SL). Together, these results demonstrate telomeric 8oxoG activates p53 and p21 to drive premature senescence.

### Acute Telomeric 8oxoG Formation Causes Telomere Fragility But Not Shortening

To determine the mechanism of telomeric 8oxoG senescence induction, we examined several telomeric endpoints. Various studies have shown cells under long-term or chronic oxidative stress exhibit accelerated telomere shortening and increased senescence (Barnes et al., 2019). Importantly, we observed premature senescence only 4 days after treatment, a timeframe in which significant telomere shortening is typically not expected or observed, particularly in telomerase proficient cells (normally occurs on the scale of weeks). We assayed for telomere shortening using Southern blot analysis of telomere restriction fragments (TRF), and found no change in the bulk telomere length 4 days after dye and light treatment (Figure S4A). However, a few critically short dysfunctional telomeres per cell (≤ 2kb) are sufficient to promote senescence in human fibroblasts (Kaul et al., 2012). Therefore, using the Telomere Shortest Length Assay (TeSLA) assay we amplified individual telomeres from cells and then performed Southern blotting to visualize the shortest telomeres. We readily detected individual telomeres much shorter than those observed in the bulk TRF population (Figure S4B). However, dye and light treatment did not increase the percent of critically short or truncated telomeres, demonstrating an acute induction of 8oxoG at telomeres does not accelerate shortening.

We next examined telomere integrity by performing telomere FISH on metaphase chromosome spreads. We used the p53ko cells to ensure damaged cells could progress from G2 to mitosis, allowing us to recover chromosomes in metaphase for analysis. Chromatid ends were scored for telomeric PNA staining as having one telomeric foci (normal), multiple foci (fragile) or no staining (signal free end) (Figure 4). In agreement with the TeSLA data, we observed little to no change in the number of signal free ends representing telomere losses or undetectable critically short telomeres, or in dicentric chromosomes representing chromosome fusions (two centromeres per chromosome) (Figures 4C-D, S4C). Consistent with a lack of 8oxoG induced telomere losses in the p53ko cells, we observed no reduction in telomere foci per nucleus 4 days after dye and light treatment in wild type interphase cells (Figure S4D). Fragile telomeres are associated with telomere replication stress (Sfeir et al., 2009; Suram et al., 2012). In both BJ and RPE FAP-TRF1 cells, we observed significant increases in fragile telomeres 24 hours after 5 minutes of dye and light treatment, suggesting telomeric 8oxoG induces replication stress (Figure 4E, F).

**Figure 4.**
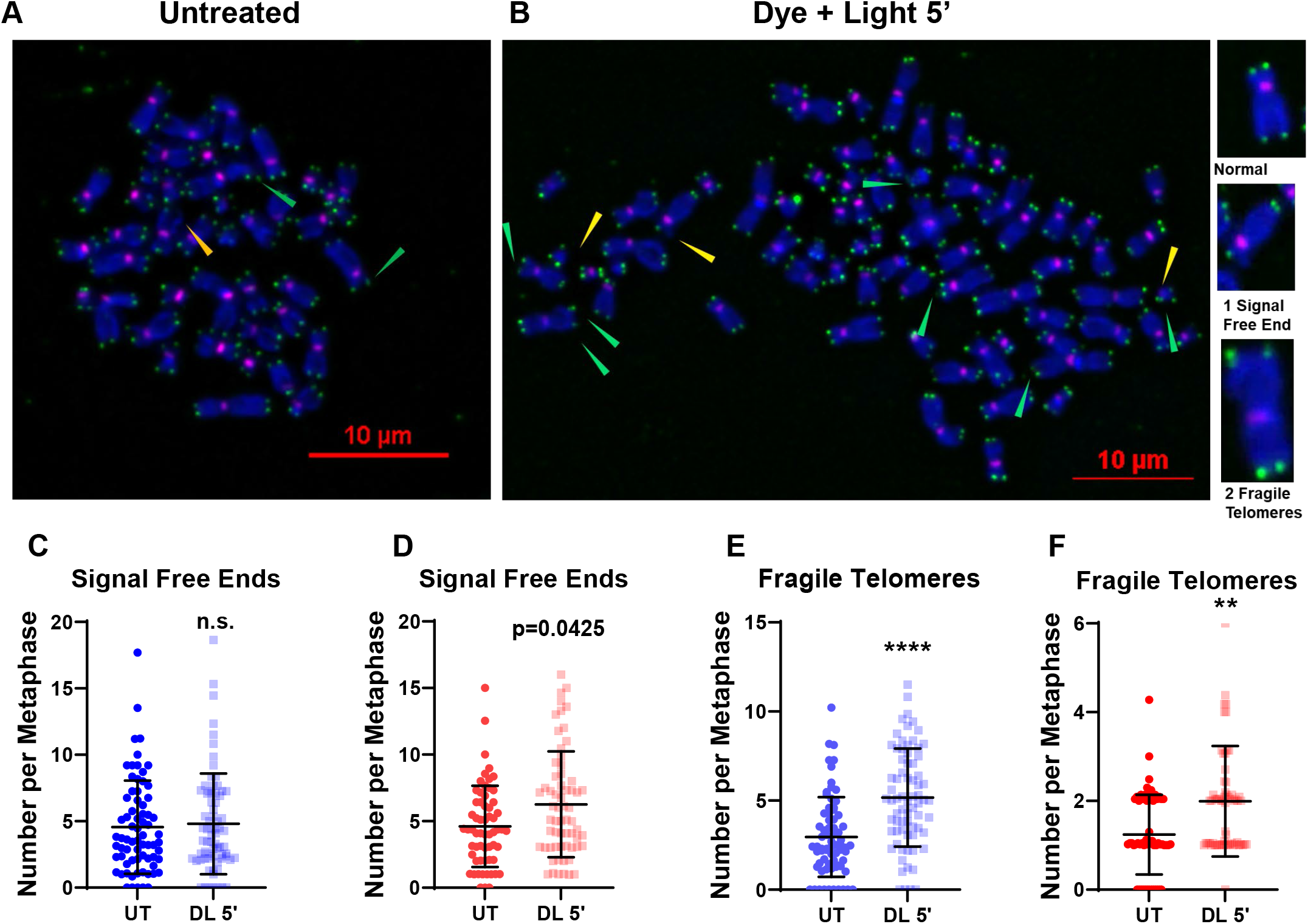
Acute Telomeric 8oxoG Formation Causes Telomere Fragility But Not Shortening. (A-B) Representative images of telo-FISH staining of metaphase chromosomes from BJ FAP-TRF1 p53 ko cells 24 h after no treatment (A) or 5 min dye + light treatment (B). Images were scored for telomeric signal free ends (yellow arrow heads) and fragile telomeres (green arrow heads). Green foci are telomeres and pink are CENPB centromeres. (C-D) Quantification of number of telomeric signal free chromatid ends per metaphase in BJ (C) and RPE (D) FAP-TRF1 cells. (E-F) Quantification of the number fragile telomeres per metaphase in BJ (E) and RPE (F) FAP-TRF1 cells. For C-F, data are the mean ± SD of at least 60 metaphases, normalized to the chromosome number. Statistical analysis was Unpaired t-Test. **p<0.01, ****p<0.0001.

Previous studies indicate that 8oxoG can disrupt TRF1 and TRF2 binding (Opresko et al., 2005). Since shelterin disruption can activate ATM and induce senescence by causing telomere deprotection, and since TRF1 loss causes telomere fragility (Denchi and de Lange, 2007; Sfeir et al., 2009), we examined whether dye and light treatment reduced TRF1 or TRF2 at telomeres. We observed no loss of FAP-mCER-TRF1 fusion protein at telomeres immediately after dye and light, as evidenced by mCerulean foci number and signal intensity at telomeres (Figure S1D and S4E). Furthermore, dye and light treatment did not significantly decrease the TRF2 signal intensity at telomeres as observed by IF-FISH (Figure S4F-G). Therefore, telomeric 8oxoG formation did not significantly alter shelterin localization at telomeres. Together these data confirm telomeric 8oxoG induces premature senescence in the absence of accelerated telomere shortening or losses, and deprotection via shelterin disruption, but instead show that 8oxoG induces telomere fragility.

### Telomeric 8oxoG Promotes Localized DDR

In addition to telomere fragility, TRF1 deletion in mouse cells significantly increases DDR positive telomeres, indicative of telomere dysfunction. These cells also exhibit growth arrest and SA-β-gal staining, which can be rescued by expressing SV40 T-antigen or p53 knockdown, both of which disrupt cell-cycle checkpoints (Martinez et al., 2009; Sfeir et al., 2009). These phenotypes are strikingly similar to what we observe with induction of telomeric 8oxoG. To determine if 8oxoG induces a DDR at telomeres, we performed IF-FISH on wild-type interphase cells 24 hours after treatment and evaluated recruitment of γH2AX and 53BP1 (Figure 5). In both BJ and RPE FAP-TRF1 cells, we observed significant increases in the percent of cells with one or more DDR positive telomeres (out of total cells), and a dramatic ∼10-fold increase in cells showing telomeres co-localized with both DDR markers compared to untreated cells (Figures 5B and 5C). We also binned this data for cells showing 0, 1-3, or 4 or more DDR positive telomeres, and found a single induction of telomeric 8oxoG significantly increases cells displaying 1-3 or ≥4 DDR positive telomeres (Figure S5A, 5SB). Previous immunofluorescence studies on metaphase chromosomes (meta-TIF) showed 4-5 γH2AX positive telomeres is predictive of replicative senescence in human fibroblasts (Kaul et al., 2012). Examining the sum of telomeres positive for either γH2AX or 53BP1 showed that 20-30% of dye and light treated cells had 4 or more DDR positive telomeres, compared to only 2% of untreated cells (Figure S5C). These results demonstrate a robust DDR at telomeres following formation of 8oxoG. We corroborated the telomeric DDR analysis in interphase cells with the meta-TIF assay by γH2AX staining of metaphase chromosome spreads 24 hours after treatment (Figure S5D). The average number of chromatid ends staining positive for both γH2AX and telomere PNA was 4.4 per metaphase, and positive for γH2AX but negative for telomere PNA was 0.9 per metaphase (Figure S5E). This confirms the majority of 8oxoG induced DDR signaling at chromatid ends was not due to telomere loss, but rather due to telomere aberrations. We also analyzed the distribution of γH2AX foci on metaphase chromosome for localization to chromatid ends versus internal sites, since it is possible interphase FISH can miss some telomeres (Cesare and Karlseder, 2012). We found that while 60% of γH2AX foci localized to chromatid ends in untreated cells, consistent with telomeres being hotspots for damage and replication stress, this increased to 83% in dye and light treated cells confirming the specificity of telomeric 8oxoG for inducing DDR at telomeres.

**Figure 5.**
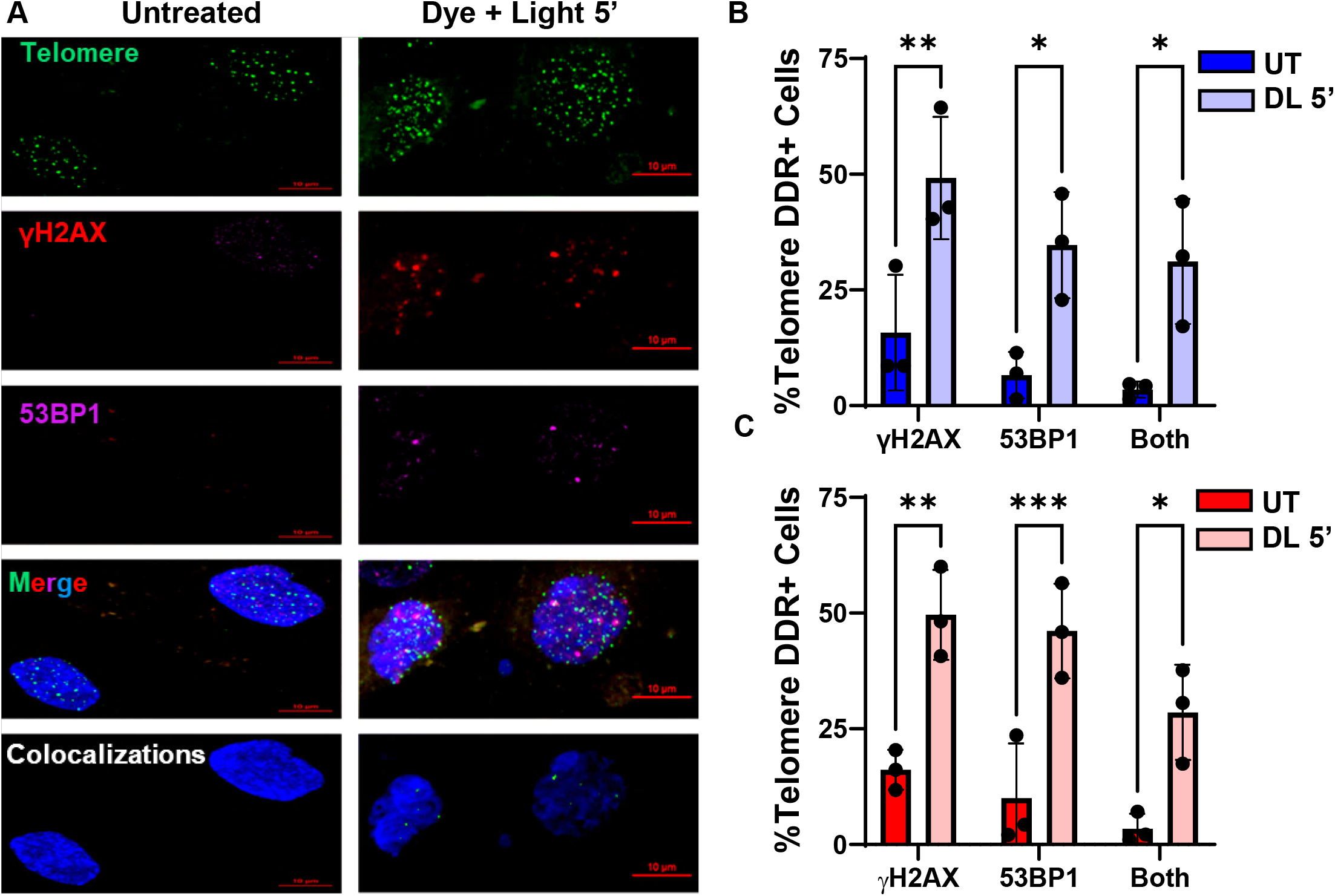
Telomeric 8oxoG Promotes a Localized DDR. (A) Representative IF images showing γH2AX (red) and 53BP1 (purple) staining with telomeres (green) by telo-FISH for BJ FAP-TRF1 cells 24 h after no treatment or 5 min dye + light. Colocalizations panel shows NIS Elements defined intersections between 53BP1 and or γH2AX with telomeres. (B-C) Quantification of percent cells exhibiting telomere foci co-localized with γH2AX, 53BP1 or both for BJ (B) and for RPE (C) FAP-TRF1 cells 24 h after no treatment or 5 min dye + light (DL 5’). Data are mean ± SD from three independent experiments in which more than 50 nuclei were analyzed per condition for each experiment. *p<0.05, **p<0.01, ***p<0.001, Two-way ANOVA.

### Replicating Cells Display a Greater DDR After Telomere 8oxoG Formation

Fragile telomeres observed after TRF1 deletion were shown to emanate from DNA replication fork stalling at telomeres, and are increased by drugs that induce replication stress such as aphidicolin (Aph) or by overexpressing oncogenes (Sfeir et al., 2009; Suram et al., 2012). To determine if S-phase cells are more sensitive to telomeric 8oxoG damage, we pre-labeled replicating cells with EdU prior to treating them with dye and light for 5 min, and then fixed them immediately after damage induction (Figure 6A). Induction of telomeric 8oxoG increased the number of γH2AX foci specifically in BJ and RPE FAP-TRF1 cells that were replicating at the time of treatment (Figure 6B-C). Consistent with this, we observed activation of the ATR/Chk1 replication stress response immediately after induction of telomeric 8oxoG (Figure S6A). In a time-course experiment we observed a significant increase in overall nuclear γH2AX signal intensity in EdU+, but not EdU-, cells one hour after treatment (Figure S6B and S6C). This signal decreased 3-12 hours after treatment, but increased again at 24 hours. This is similar to reported observations in hydrogen peroxide treated cells in which the second wave of DDR is proposed to result from increased replication fork encounters with DNA lesions or repair intermediates (Venkatachalam et al., 2017). Importantly, we also observed this trend of an immediate DDR activation, transient drop, and rebound at 24 hours at telomeres (Figure S6D-F).

**Figure 6.**
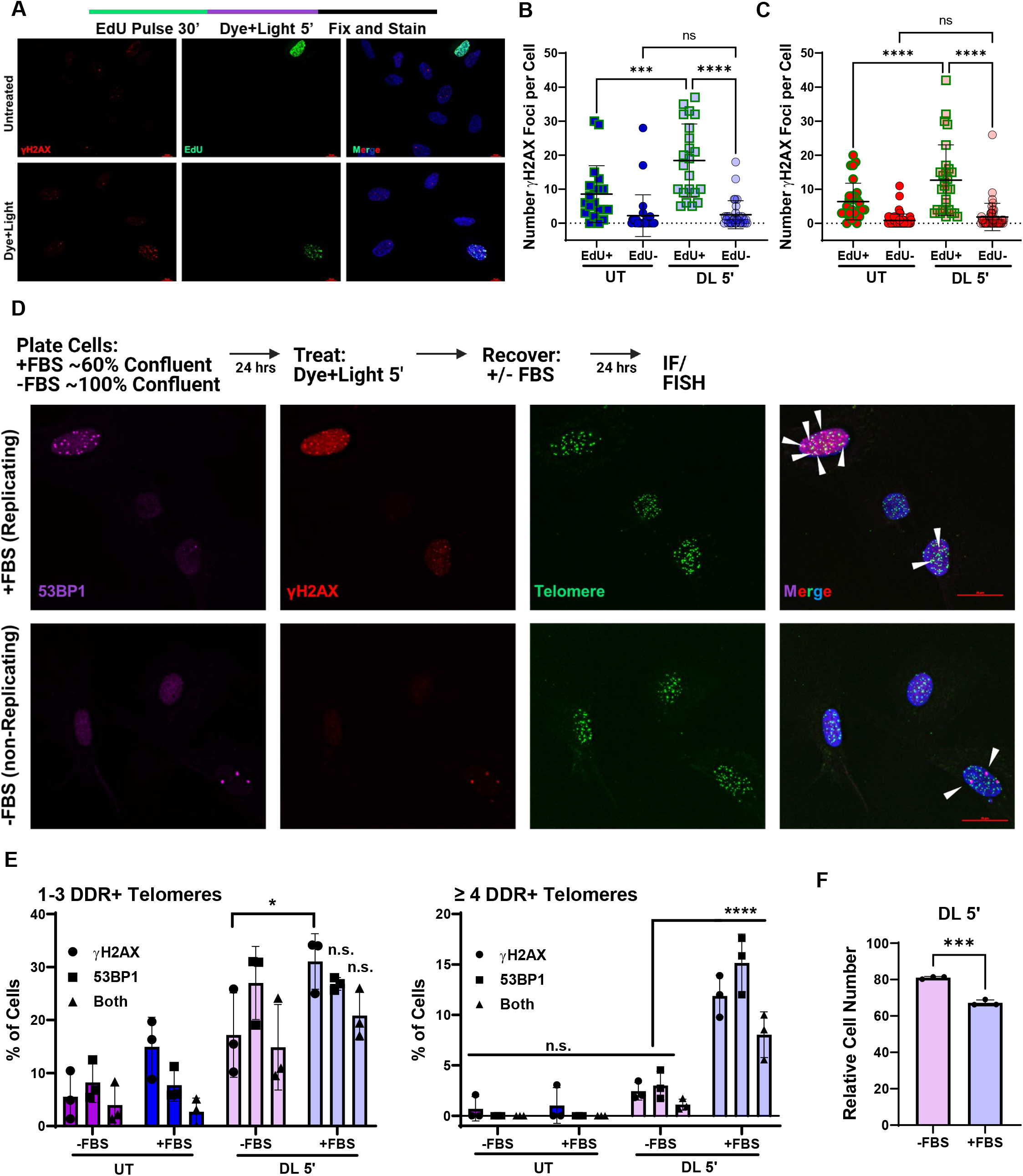
Replicating Cells Display a Greater DDR After Telomere 8oxoG Formation. (A) Schematic shows experiment for EdU labelling of S-phase cells. Representative IF image of γH2AX (red) and EdU (green) staining of BJ FAP-TRF1 cells after 0 h recovery from 5 min dye + light. (B, C) Quantification of number of γH2AX foci per EdU negative (-) and EdU positive (+) cells for BJ (B) and RPE (C) FAP-TRF1 cells. For each condition, over 60 nuclei were analyzed and scored as EdU + or -. (D) Schematic of experiment for telomere DDR imaging in replicating (cells grown with 10% FBS; +FBS) and non-replicating (cells grown with 0.1% FBS; -FBS) BJ FAP-TRF1 cells. Representative IF/FISH images shown below. (E) Quantification of the percent of cells with 1-3 or ≥4 DDR+ (γH2AX, 53BP1, or both) telomeres as in D. Data are the averages of 3 experiments each examining > 70 nuclei per condition. (F) Cells were seeded in media with 0.1% (-) of 10% (+) FBS, treated the next day with 5 min dye + light, and then recovered 24 h with 0.1 (-) or 10% (+) FBS. All cells were cultured in 10% FBS media another 4 days before counting. All data are mean ± SD, and were analyzed by One-way ANOVA (B, C), Two-way ANOVA (E), or t-test (F). *p<0.05, ***p<0.001, ****p<0.0001.

To investigate a potential role for DNA replication in telomere 8oxoG induced DDR and senescence, we examined quiescent cells. Fibroblasts quickly synchronize to G0/G1 when serum starved and confluent (Krek and DeCaprio, 1995). Therefore, we seeded BJ cells at nearly 100% confluence with only 0.1% FBS. 24 hours after seeding, cells were treated with dye and 5 min light, and then recovered for 24 hours with 0.1% FBS media (Figure 6D). Control cells were seeded and grown in 10% FBS as normal, treated, and recovered in media with 10% FBS. We confirmed quiescence by examining EdU incorporation and cyclin A expression (Figure S6G-I). 24 hours after induction of telomeric 8oxoG, replicating (+FBS) cells showed an increase both in cells with 1-3 DDR+ telomeres, and 4 or more DDR+ telomeres (Figure 6E), as observed previously (Figures S5A-B). However, while cells grown in 0.1% FBS showed an increase in 1-3 DDR+ telomeres per cell, there was no significant increase in cells with 4 or more DDR+ telomeres. Because 4 or more DDR+ telomeres is predictive of senescence (Kaul et al., 2012), we tested if preventing DNA replication for 24 hours after dye and light treatment would rescue growth arrest. Indeed, quiescent cells treated and recovered in 0.1% FBS media, before allowing to grow in 10% FBS media, displayed improved growth compared to replicating cells treated and recovered in 10% FBS media (Figure 6F). These findings show DNA replication following formation of 8oxoG at telomeres, is a major contributor to the resulting DDR activation and growth reduction.

### 8oxoG Directly Disrupts Telomere Replication

To directly interrogate replication stress at telomeres, we tested for mitotic DNA synthesis (MiDAS). MiDAS is a DNA repair pathway activated at difficult to replicate regions to enable completion of DNA synthesis, and is detected by EdU incorporation during mitosis (Minocherhomji et al., 2015). We treated p53ko RPE FAP-TRF1 cells with dye and light and allowed them to recover 16 hours, before adding the CDK1i R0-3306 for 24 hours, to arrest cells at the G2/M boundary. Aph treatment was used as a positive control (Figure S7A). Cells were then released into media containing EdU and colcemid, allowing for DNA synthesis, and halting cells in metaphase (Figure 7A). As expected, in Aph treated cells we observed robust EdU incorporation at mostly internal chromosomal regions, which was often coincident with chromosome breaks (Figure S7A). In contrast, dye and light treated cells showed primarily telomeric EdU foci (Figure S7B). At least one telomere MiDAS event occurred in 79% of dye and light treated cells, compared to only 39% of untreated cells.

**Figure 7.**
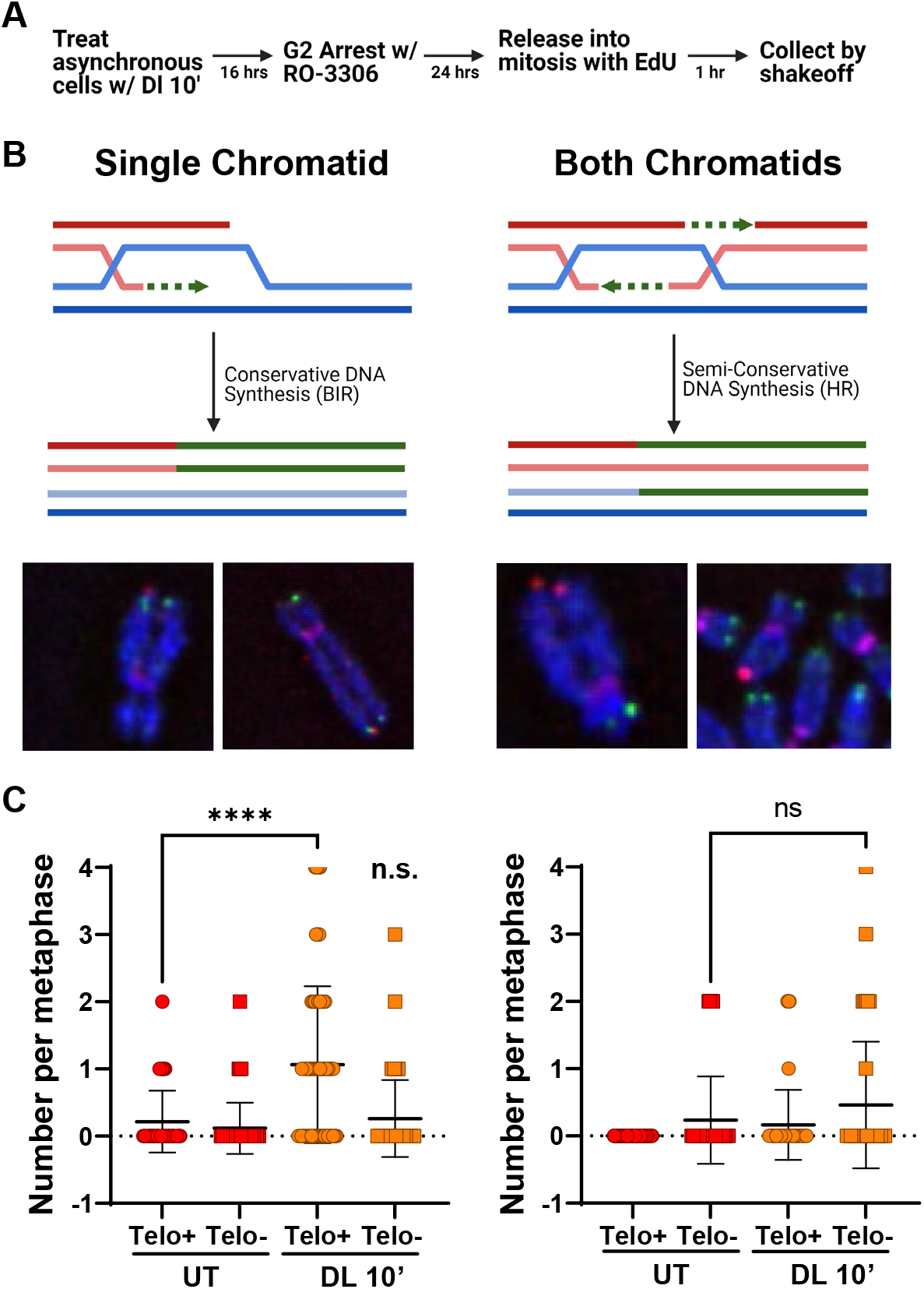
8oxoG Directly Disrupts Telomere Replication. (A) Schematic of MiDAS experiment in p53ko RPE FAP-TRF1 cells. (B) Schematics for EdU events at a single chromatid (BIR) and both chromatids (HR). Representative images from dye and light treated RPE FAP-TRF1 cells are shown below. (C) Quantification of telomere MiDAS events at a single chromatid (left) or both chromatids (right). Events are scored for as chromatid end staining positive (Telo+) or negative (Telo-) for telomeric PNA. Data are from > 50 metaphases from two independent experiments. One-way ANOVA, ****p<0.0001.

Previous studies have shown telomere MiDAS primarily occurs in a conservative DNA synthesis manner on a single chromatid, consistent with break-induced-replication (BIR), in contrast to homologous recombination (HR) which requires semi-conservative synthesis on both chromatids (Figure 7B) (Min et al., 2017; Özer et al., 2018). Therefore, we scored MiDAS events as occurring on a single chromatid or both chromatids, and whether the chromatid end was positive for telomere PNA staining. While untreated cells displayed an average of 0.2-0.3 telomere MiDAS events per metaphase (single or both; median 0), dye and light treated cells showed a significant increase in single chromatid telomere MiDAS, with an average and median number of 1 per metaphase (Figure 7C). These single chromatid events almost exclusively occurred on telomere PNA positive chromatid ends, consistent with BIR. While not significant, there was a slight increase in MiDAS on both chromatids following dye and light treatment on chromatids negative for telomere PNA, which is consistent with HR. These data indicate that acute telomeric 8oxoG formation leads to mitotic DNA synthesis, suggesting the lesions prevented the completion on telomeric DNA replication in S-phase.

## DISCUSSION

A wealth of evidence indicates that oxidative stress both enhances cellular aging and accelerates telomere dysfunction (Barnes et al., 2019), and here we demonstrate a direct causal link between these two ROS-induced cellular outcomes. Oxidative stress had been proposed to hasten telomere shortening and the onset of senescence by producing 8oxoG lesions in highly susceptible TTAGGG repeats (von Zglinicki, 2002). Whether telomeric 8oxoG has a causal role in driving senescence could not be tested previously, because telomeres comprise a tiny fraction of the genome, and oxidants used to produce 8oxoG modify numerous cellular components and alter redox signaling. We overcame these barriers by using a chemoptogenetic precision tool that induces singlet oxygen mediated 8oxoG formation exclusively at telomeres. We demonstrate acute telomeric 8oxoG formation is sufficient to trigger rapid premature senescence in the absence of telomere shortening or losses in primary and hTERT expressing human cells. Instead, we observed telomere fragility, DNA damage signaling, and replication stress at telomeres. Mechanistically, our data are consistent with a model in which 8oxoG itself, and/or subsequent repair intermediates, stall DNA replication at the telomeres, leading to a robust induction of p53 signaling to arrest cell growth and enforce premature senescence.

Here, we found formation of 8oxoG exclusively at telomeres induces multiple hallmarks of premature senescence in as few as four days. We confirmed increases in SA-β-gal activity, CCFs, and mitochondrial activity as well as reductions in colony formation, LaminB1 expression, EdU incorporation, and RB phosphorylation following induction of telomeric 8oxoG, phenotypes previously observed in replicative senescence, or oncogene and DNA damaged induced premature senescence (Hernandez-Segura et al., 2018). Several of these phenotypes were rescued when ATM was inhibited pharmacologically or p53 was deleted genetically, consistent with other models of premature senescence (Beausejour, 2003; Kang et al., 2017; Kuk et al., 2019). The rapid timescale of telomeric 8oxoG induced senescence would not typically allow for significant telomere shortening, which we indeed confirmed. Notably, in HeLa FAP-TRF1 cells we only observed telomere shortening and losses after chronic 8oxoG formation in wild-type and especially repair deficient cells (Fouquerel et al., 2019), which raises the possibility that chronic damage may also accelerate shortening in non-diseased cells that continue to proliferate. Never-the-less, our data demonstrate that telomeres have a pronounced sensitivity to the presence of oxidative stress induced 8oxoG, but that this is independent of shortening. Instead, we propose the sensitivity results from DNA replication slowing or stalling, resulting in a robust DDR.

Telomeres exist in a “t-loop” structure when organized by shelterin to prevent erroneous recognition as DSBs and the DDR. Additionally, shelterin proteins can directly inhibit the activities of the HR and end-joining pathways from acting at telomeres even when the t-loop structure is not present but DDR is activated (intermediate-state), thereby preventing fusions (Cesare and Karlseder, 2012; Van Ly et al., 2018). These observations, together with our results, highlight how readily damaged telomeres can be sensed by the DDR and suggest that 8oxoG may promote an intermediate-state. We found that while telomeric 8oxoG formation increased γH2AX and 53BP1 recruitment to telomeres within 24 hours, it did not increase telomere losses and truncated telomeres which could arise from direct breaks, or chromosome fusions which could reflect TRF2 loss. Instead of telomere shortening and losses, our data suggest acute 8oxoG disrupts DNA replication at telomeres, perhaps generating an intermediate-state and DDR activation, which promotes senescence in cells with intact p53 signaling pathways. Replication stress is defined as the slowing or stalling of DNA replication forks, and leads to robust increases in telomere fragility (Sfeir et al., 2009; Suram et al., 2012). While still structurally undefined, fragile telomeres are believed to represent unreplicated regions in the telomere causing altered chromatinization (Ruis and Boulton, 2021). We found a single induction of telomeric 8oxoG enhances telomere fragility. While acute telomere 8oxoG also induces fragility in HeLa cells, this increase was not significant compared to repair deficient cells (Fouquerel et al., 2019), suggesting non-diseased cells are more sensitive than cancer cells.

When barriers prevent the completion of replication, cells can activate the mitotic DNA synthesis (MiDAS) pathway to resume synthesis before cell division. Following induction of telomeric 8oxoG, we observe a robust increase in single chromatid MiDAS events, which is consistent with other models of telomere replication stress (Min et al., 2017; Özer et al., 2018; Yang et al., 2020). We verified the telomere DDR response we observed was due to DNA replication by comparing replicating cells with serum-starved quiescent cells. While the serum starved cells had an increase in telomere DDR, it was significantly weaker than in replicating cells. Specifically, serum starved cells did not show a large increase in the percentage of cells with 4 or more DDR+ telomeres, a phenotype previously shown to correlate with replicative senescence (Kaul et al., 2012). Together, these data along with the telomere fragility and Chk1 phosphorylation demonstrate the formation of 8oxoG at telomeres leads to replication stress.

How does 8oxoG impact replication? While studies have largely focused on the mutagenic consequences of 8oxoG, we argue mutagenesis is unlikely a primary contributor to the cellular phenotypic outcomes, since DDR foci arose immediately after induction of the lesion. Our data suggest 8oxoG stalls replication at the telomeres. *In vitro* studies examining replicative DNA polymerase delta (Pol δ) synthesis across 8oxoG in the template have shown this lesion is not as strong of a block as bulky lesions caused by UV light or cisplatin. However, Pol δ stalling at the 8oxoG lesion has been reported, especially when incorporating C, even in the presence of its accessory factors RFC and PCNA (Markkanen et al., 2012). Further support for Pol δ stalling derives from evidence that translesion polymerases η and λ function in bypass of 8oxoG (Markkanen et al., 2012; McCulloch et al., 2009). Importantly, many of the polymerase biochemical studies were conducted using dNTP concentrations above cellular relevant concentrations, on non-telomeric templates (Haracska et al., 2002; Schmitt et al., 2009). The human mitochondrial replisome shows substantial stalling at 8oxoG in reactions containing cellular dNTP levels (Stojkovic et al., 2016), suggesting previous biochemical studies may have underestimated the impact of 8oxoG on replication fork progression in cells. Furthermore, since difficult-to-replicate sequences, such as telomeres, themselves can impede Pol δ upon replication stress, future biochemical studies are warranted to study Pol δ telomere synthesis in the presence of physiological dNTP levels and 8oxoG within telomere templates.

The key finding from our study, that a minor oxidative base lesion arising within the telomeres is sufficient to induce premature senescence in the absence of telomere shortening, is surprising, but provides a mechanistic explanation for studies of telomere dysfunction arising *in vivo* in various contexts (Victorelli and Passos, 2017). Cardiomyocytes from mice, and hippocampal neurons and hepatocytes in baboon, show increased DDR positive telomeres with age, with no appreciable shortening despite the presence of senescence markers (Anderson et al., 2019; Fumagalli et al., 2012). Oxidative stress is implicated in generating DDR positive telomeres in liver and intestinal cells, also without shortening, in mouse models of liver damage and chronic low-grade inflammation, respectively (Jurk et al., 2014; Lagnado et al., 2021). Human melanocytic nevi also senesce in the absence of telomere shortening (Michaloglou et al., 2005). Moreover, in cell culture models of replicative senescence and ionizing radiation or hydrogen peroxide induced premature senescence, DDR foci persist or accumulate at telomeres, long after their detection at non-telomere sites, irrespective of telomere length (Fumagalli et al., 2012; Hewitt et al., 2012; Kaul et al., 2012). Together these reports demonstrate cells can senesce independent of telomere attrition, however telomeres in these cells are largely positive for DDR foci. Our data showing that 8oxoG does not induce telomere shortening in an acute treatment, but significantly elevates their DDR signaling, provides a possible mechanism. Consistent with previous work (Kaul et al., 2012), we also observed the vast majority of γH2AX foci at chromatid ends occurred at those positive for telomere PNA staining, indicating the DDR was not due to telomere loss.

Importantly, our study demonstrates premature senescence in primary and non-diseased human cells following the induction of a common, physiological oxidative DNA lesion targeted to the telomere. Oxidative stress is a ubiquitous source of DNA damage humans experience due to endogenous metabolism and inflammation, exogenous environmental sources, and life-stress and epidemiological studies have shown an increase in 8oxoG levels in aged humans (Gan et al., 2018; Osterod et al., 2001). Our results highlight the importance of understanding how and where this DNA lesion arises within human genomes, since its presence at telomeres alone is sufficient to rapidly advance cellular aging. Thus, these studies reveal a novel mechanism of telomere driven senescence linked to oxidative stress.

## ACKNOWLEDGEMENTS

This work was supported by NIH grants F32AG067710-01 (to R.P.B.), R35ES030396 and R01CA207342 (to P.L.O), and R01EB017268 (to M.P.B.). This project used the UPMC Hillman Cancer Center CF that is supported in part by award P30CA047904. This project was also supported by the UPMC Hillman Cancer Center Postdoctoral Fellowship for Innovative Cancer Research (R.P.B).

## AUTHOR CONTRIBUTIONS

R.P.B. and P.L.O conceived of the study. R.P.B. and P.L.O designed the experiments. R.P.B. performed most of the experiments, except M.D.R. conducted RPE metaphase spread analysis and assisted with some Beta-gal experiments, S.A.T. conducted qPCR experiments, A.C.D. performed TESLA experiments, V.R. performed Seahorse analysis, and J.S.O. conducted the RNA-Seq analyses. M.P.B. provided the MG2I dye and 660 nm LED irradiator. R.P.B. and P.L.O. wrote the manuscript with assistance from the other authors.

## SUPPLEMENTAL INFORMATION

**Figure S1. Acute Telomeric 8oxoG Initiates Rapid Premature Senescence in Non-Diseased Cells, Related to Figure 1**.

(A) Representative images of YFP-XRCC1 (red) localization to telomeres FAP-mCer-TRF1 (cyan) after 10 min dye + light treatment, obtained by direct mCer and YFP visualization.

(B) Quantification of percent YFP-XRCC1 positive telomeres per nuclei after no treatment (UT) or 10 min dye + light treatment in wild-type or OGG1 knock-out (ko) RPE FAP-TRF1 cells. N ≥ 25.

(C) Quantification of YFP-XRCC1 signal intensity at telomeric foci as shown in (A). Box plots represent quantification of data from at least 935 foci per condition.

(D) Quantification of number (#) of FAP-mCer-TRF1 foci per cell by direct mCer visualization of untreated cells (UT) or 10 min after dye + light (DL 10’).

(E) Quantification of percent YFP-XRCC1 positive telomeres per nuclei after 10 min dye + light exposure with pretreatment of 100 µM NaN_3_ for 15 min prior to light exposure (NaAz) or with no pretreatment (-).

(F) Immunoblot for FAP-TRF1 and OGG1 in whole cell extracts from RPE FAP-TRF1 wild type and OGG1 ko cells. Actin shown as loading control. * indicates non-specific band stained by anti-OGG1.

(G) Cell counts of RPE FAP-TRF1 clone 18 obtained 4 days after recovery from indicated treatments.

(H) Cell counts of RPE FAP-TRF1 (red) and BJ FAP-TRF1 (blue) cells 4 days after recovery from 1, 5, 10, or 20 min dye + light treatments relative to untreated.

(I-J) Cell counts of parental BJ-hTERT (I) or RPE-hTERT (J) cells obtained 4 days after recovery from indicated treatments.

(K) Cell counts of FAP-TRF1 expressing HeLa and U2OS clones obtained 4 days after recovery from 5 min dye + light treatment relative to untreated.

(L) Representative image of FAP-mCER-TRF1 protein colocalization with telomeres in bulk population primary BJ cells visualized by mCER IF (pink) with telo-FISH (green).

(M-N) Cell counts of BJ FAP-TRF1 (M) or RPE FAP-TRF1 (N) cells 4 days after recovery from 1-hour treatments with 2.5 or 10 mM KBrO_3_, and for BJ FAP-TRF1 with 50 μM etoposide (ETP).

(O) Cell counts of BJ (E) or RPE (F) FAP-TRF1 cells obtained 24 hours after recovery from 5 min dye + light treatment relative to untreated.

(P) Flow cytometry plots of RPE FAP-TRF1 cells showing gating based on EdU and propidium iodine staining for various cell cycle phases 24 h after no treatment or exposure to dye, light, dye + 5’ light, 20 J/m^2^ UVC, or 1 hour treatment with 2.5 or 10 mM KBrO_3_.

(Q) Representative images of β-galactosidase positive BJ FAP-TRF1 cells obtained 4 days after recovery from indicated treatments (From Figure 1H-I). All graphs show the mean ± SD. B-C and analyzed by One-way ANOVA. *p<0.05, **p<0.01, ***p<0.001, ****p<0.0001.

**Figure S2. Telomeric 8oxoG Production Increases Cytoplasmic DNA, Related to Figure 2**.

(A) Representative image of Lamin B1 and Lamin A/C IF staining of BJ FAP-TRF1 cells obtained 4 days after recovery from 5 min dye + light. Gold box is zoom shown in Figure 2G.

(B) Immunoblot of Lamin A/C and Lamin B1 in BJ FAP-TRF1 cells obtained 9 days after recovery from indicated treatments.

(C) Size of nuclear area (μm^2^) of BJ FAP-TRF1 (blue) or RPE FAP-TRF1 (red) cells obtained 4 days after recovery from no treatment or 5 min dye + light. At least 80 nuclei examined per condition.

(D) Representative scatterplots of Annexin V (y-axis) and propidium iodine (x-axis) staining of cells 4 days after the indicated treatments.

(E) PFGE of cells in agarose plugs and SybrGreen staining of genomic DNA from BJ FAP-TRF1 and RPE FAP-TRF1 cells untreated (UT) or after 0 or 24 h recovery from 5 min dye + light (D+L) treatment. Treatments for 1 hour with 1 or 10 mM H_2_O_2_, or 40 mM KBrO_3_, used as positive controls.

Panels C and D are means ± SD and analyzed by t-test. ***p<0.001, ****p<0.0001.

**Figure S3. p53 DNA Damage Signaling is Required for 8oxoG Induced Senescence, Related to Figure 3**

(A) Immunoblot of phosphorylated Rb from BJ FAP-TRF1 cells 4 days recovery after indicated treatments.

(B, C) Immunoblots of BJ (C) and RPE (D) FAP-TRF1 cells 3 hours after 5 min dye + light treatment. Cells were also pre-and post-treated with ATMi KU55933 (10 μM) or KU60019 (1 μM) or DMSO solvent control. * indicates non-specific band.

(D, E) Volcano plots of gene expression changes in RPE (D) and BJ (E) FAP-TRF1 cells 24 hours after dye + light treatment. Each dot is a gene and red dots are significantly up or down-regulated. HeLa cells were analyzed the same, but no significant changes were observed.

(F, G) Analysis of gene expression changes in RPE (F) and BJ (G) FAP-TRF1 cells 24 hours after dye + light treatment, as a function of chromosome position. Each dot is a 10kb bin, and the red line is the average.

(H) qPCR analysis of p16 mRNA (*CDK2NA*) in BJ FAP-TRF1 cells, 4 days after the indicated treatments. Data were analyzed by One-Way ANOVA.

(I) Cell counts of wild-type and p53ko RPE FAP-TRF1 cells 4 days after treatment with dye + light for the indicated times.

(J) Cell counts of wild-type and p53ko RPE and BJ FAP-TRF1 cells 4 days after treatment with 10 mM (RPE) or 2.5 mM (BJ) KBrO_3_.

(K, L) 23 hours after treatment, wild-type and p53ko RPE FAP-TRF1 (K) and BJ FAP-TRF1 (L) cells were pulsed with EdU for 1 hour, and then analyzed by microscopy for p21 and EdU staining. In each condition, cells were categorized as EdU + or – populations, and the nuclear p21 signal intensity was graphed. Representative IF images are also shown.

For panels H-L, data are the mean ± SD and analyzed by One or Two-way ANOVA. *p<0.05, **p<0.01, ***p<0.001, ****p<0.0001.

**Figure S4. Acute Telomeric 8oxoG Formation Causes Telomere Fragility But Not Shortening, Related to Figure 4**

(A) Representative Southern blot for telomere restriction fragment length analysis obtained from BJ and RPE FAP-TRF1 after 4 days recovery from no treatment (UT) or dye alone, or 5 min light alone, or dye + light together.

(B) Representative TeSLA Southern blot obtained from BJ and RPE FAP-TRF1 after 4 days recovery from no treatment (UT) or 5 min dye + light. Each lane is an independent PCR from the same pool of genomic DNA.

(C) Quantification of dicentric chromosomes defined as two centromeric foci for p53 ko BJ (blue) and RPE (red) FAP-TRF1 cells 24 h after 5 min dye + light treatment.

(D) Quantification of telomere foci per cell as measured by telo-FISH for BJ (blue) and RPE (red) FAP-TRF1 cells 4 days after no treatment (UT) or 5 min dye + light. Bars represent the means ± SD from 80 to 184 nuclei per condition. Non-significant (One-way ANOVA).

(E) Quantification of mCER signal intensity per telomere foci from FAP-mCER-TRF1 in wild-type RPE FAP-TRF1 cells after no treatment (UT) or 10 min dye + light (DL 10’). Box plot shows means from 1000 to 1300 nuclei per condition. Non-significant (One-way ANOVA).

(F) Representative IF images for TRF2 (pink) colocalized with telomeres by telo-FISH (green) in RPE FAP-TRF1 cells immediately after 5’ dye + light treatment. DNA stained with DAPI.

(G) Quantification of TRF2 signal intensity per telomere foci in RPE FAP-TRF1 (red) and BJ FAP-TRF1 (blue) cells from (F). Box plot shows medians (bar) and means (+) from 1460 to 3499 nuclei per condition. Non-significant (One-way ANOVA). *p<0.05, **p<0.01, ***p<0.001, ****p<0.0001.

**Figure S5. Telomeric 8oxoG Promotes a Localized DDR, Related to Figure 5**

(A-B) Quantification of the percent of cells showing 0, 1-3 or ≥4 telomeric foci co-localized with γH2AX, 53BP1 or both for BJ FAP-TRF1 cells (A) and for RPE FAP-TRF1 cells (B) 24 hours after no treatment or 5 min dye + light treatment

(C) The % of cells with ≥4 γH2AX or 53BP1 positive telomeres from each experiment in Figure 5 is shown, and summed. The yellow cell is the average of all 3 experiments.

(D) Representative image of meta-TIF chromosome spread from RPE FAP-TRF1 cell 24 hours after a 5 min dye + light treatment stained for γH2AX (red), telomere PNA (green) and for DNA by DAPI (blue).

(E) Quantification from meta-TIF assay of γH2AX positive chromatid ends lacking telomere staining (Telo -) or co-localized with telomeric PNA (Telo +) by telo-FISH as shown in (C). N = 33 metaphases.

(F) Pie chart from meta-TIF assay shows distribution of γH2AX foci located at chromatid ends (telomeric) verses internal (non-telomeric) sites by IF and teloFISH as shown in (C). All graphs show the mean ± SD.

**Figure S6. Replicating Cells Display a Greater DDR After Telomere 8oxoG Formation, related to Figure 6**.

(A) Immunoblot of phosphorylated Chk1 (S317) and H2AX from untreated (UT) BJ FAP-TRF1 and cells treated with 5 min dye + light and recovered for the indicated times. UV = 20 J/m^2^ UVC.

(B) Representative IF image of γH2AX (red) and EdU (green) for total of 24 h recovery (23 h fresh media + 1 h EdU media).

(C) (Top) Schematic shows experiment for EdU labelling of S-phase cells of BJ FAP-TRF1 cells after 5 min dye + light treatment and total recovery time. 1 hour before harvest, cells were pulsed with EdU. (Bottom) Quantification of total nuclear γH2AX intensity as shown in (B) for EdU+ and EdU-cells for various recovery time points. Data are means from at least 100 total nuclei per condition.

(D-F) Quantification of the number of telomeres per nuclei co-localized with γH2AX, 53BP1 or both (DDR+) in BJ FAP-TRF1 cells untreated or after 5 min dye + light treatment and 0h, 3h, 24h or 4 days recovery. Data are mean ± SD from at least 60 nuclei per condition.

(G) Representative brightfield images of BJ FAP-TRF1 cells grown with the indicated FBS concentration and after the indicated treatment.

(H, I) Quantification of EdU positive cells (H) and Cyclin A nuclear signal (I) in BJ FAP-TRF1 cells 24 hours after the indicated treatments and recovery in 10% FBS (+) or 0.1% FBS (-), as in Figure 6D. In panel I significance is shown for -FBS cells relative to +FBS cells.

All graphs show the mean ± SD and were analyzed by One-way ANOVA, *p<0.05, **p<0.01, ***p<0.001, ****p<0.0001..

**Figure S7. 8oxoG Directly Disrupts Telomere Replication, Related to Figure 7**.

(A) Representative images of MiDAS assay for EdU incorporation on metaphase chromosomes of p53ko RPE FAP-TRF1 cells treated with 0.4 μM APH exposure for 16 h before adding CDK1i and additional 24-hour incubation. Images shows EdU (red) colocalization with telomeric DNA (green) and at internal non-telomeric sites. DNA stained with DAPI.

(B) Representative images of MiDAS assay for EdU incorporation on metaphase chromosomes of p53ko RPE FAP-TRF1 cells 24 hours after 5 min dye + light. Arrows point to EdU (red) co-colocalized with telomeric DNA (green).

**Supplemental Table 1**

RPE and BJ FAP-TRF1 RNA-seq data were analyzed by Hallmark gene set enrichment and the adjusted p values (padj) and normalized enrichment score (NES) for pathways with p > 0.005 are shown. Green indicated upregulation and red downregulation.

## METHOD DETAILS

### Cell Culture and Cell Line Generation

hTERT expressing BJ and RPE cells, as well as primary BJ cells were purchased from ATCC and tested for mycoplasma. BJ cells were grown in DMEM (Gibco) with 10% Hyclone FBS and 1% penicillin/streptomycin. RPE cells were grown in DMEM/F12 (Gibco) with 10% FBS (Gibco) and 1% penicillin/streptomycin. To generate FAP-mCER-TRF1 expressing clones, HEK 293T cells were transfected with pLVX-FAP-TRF1 and Mission Packaging Mix (Sigma) to produce lentivirus. hTERT BJ and RPE cells were infected with virus 48 and 72 hours post transfection, and then selected with 1 mg/ml G418 (Gibco). Surviving cells were single-cell cloned and expanded before checking for FAP-mCER-TRF1 expression. Primary BJ cells were infected and selected the same way, but were not single-cell cloned. After initial selection, FAP-TRF1 expression was maintained with 500µg/ml G418. U2OS and HeLa FAP-TRF1 cells were previously described (Fouquerel et al., 2019). Except for 293T cells, all cells are maintained at 5% O_2_.

To generate knockout cell lines, 293T cells were transfected with pLentiCRISPR V2 plasmids encoding guide RNAs to the respective targets and S. pyogenes Cas9 (GeneScript). FAP-TRF1 expressing cells were infected with lentivirus as above and selected with 1 µg/ml (BJ) or 15 µg/ml (RPE) Puromycin (Gibco). After selection and death of uninfected cells, the infected cells were expanded and expression of targeted protein(s) was determined by western blotting.

### Cell Treatments

For dye and light treatments, cells were plated at an appropriate density for the experiment overnight. The next day, cells were changed to Optimem (Gibco) and incubated at 37°C for 15 minutes before adding 100nM MG2I for another 15 minutes. Cells were then placed in the lightbox and exposed to a high intensity 660 nm LED light at 100 mW/cm^2^ for 5 min (unless indicated otherwise). KBrO_3_ and etoposide were added in Optimem at the indicated concentrations for 1 hour.

### Growth Analyses

For cell counting experiments, cells were plated at a low density in 6-well or 6-cm plates overnight. Cells were treated as indicated and returned to the incubator and recovered for the indicated amount of time (typically 4 days). Cells were detached from the plates, resuspended, and counted on a Beckman Coulter Counter. Each experiment had 2-3 technical replicates, which were averaged.

### Senescence Associated Beta-Galactosidase Assay

Detection of β-gal activity was done according to manufacturer’s instructions (Cell Signaling). Briefly, cells were washed with PBS, and then fixed at room temperature for 10 minutes. After 2 rinses with PBS, cells were incubated overnight at 37°C with X-gal staining solution with no CO_2_. Images were acquired with a Nikon brightfield microscope with DS-Fi3 camera. Images were scored in NIS Elements. At least 200 cells were counted per technical replicate.

### Colony Formation Assay

RPE FAP-TRF1 cells were plated in 6 cm plates overnight. The cells were treated with dye and light the next day and immediately detached, counted, and plated in triplicate in 6 well plates. After 7-8 days, the colonies were fixed on ice in 100% methanol, stained with crystal violet solution, and then counted manually.

### Immunofluorescence and FISH

Cells were seeded on coverslips and treated as indicated. Following treatment and/or recovery, cells were washed with PBS and fixed at room temperature with 4% formaldehyde. If cells were extracted before fixation, they were treated on ice with ice cold CSK buffer (100 mM NaCl, 3 mM MgCl_2_, 300 mM glucose, 10 mM Pipes pH 6.8, 0.5% Triton X-100, and protease inhibitors tablet (Roche). Fixed cells were rinsed with 1% BSA in PBS, and washed 3x with PBS-Triton 0.2% before blocking with 10% normal goat serum, 1% BSA, and 0.1% Triton-x. Cells were incubated overnight at 4°C with indicated primary antibodies. Next day cells were washed with PBS-T 3x before incubating with secondary antibodies and washing again 3x with PBS-T. If FISH was performed, the cells were re-fixed with 4% formaldehyde and rinsed with 1% BSA in PBS, and then dehydrated with 70%, 90%, and 100% ethanol for 5 minutes. Telomeric PNA probe was diluted 1:100 (PNABio) prepared in 70% formamide, 10mM Tris HCl pH 7.5, 1x Maleic Acid buffer, 1x MgCl_2_ buffer, and boiled for 5 minutes before returning to ice. Coverslips were then hybridized in humid chambers at room temperature for 2 hours, or overnight at 4°C. The cells were washed with 70% formamide and 10 mM tris HCl pH 7.5 2 times, PBS-T 3 times, and rinsed in water before staining with DAPI and mounting. Image acquisition was performed with a Nikon Ti inverted fluorescence microscope. Z stacks of 0.2 μm thickness were captured and images were deconvolved using NIS Elements Advance Research software algorithm. To detect EdU incorporation, Click chemistry was performed after the secondary antibody washes according to the manufacturer’s instructions (Thermo).

### Metaphase Spreads

Chromosome spreads were prepared by incubating cells with 0.05 μg/ml colcemid for 2 hours prior to harvesting with trypsin. Cells were incubated with 75mM KCl for 8 minutes at 37°C and fixed in methanol and glacial acetic acid (3:1). Cells were dropped on washed slides and dried overnight before fixation in 4% formaldehyde. Slides were treated with RNaseA and Pepsin at 37°C, and then dehydrated. FISH was performed as above, and included a CENPB (PNABio) probe in addition to the telomere probe.

### Pulsed-Field Gel Electrophoresis of Cells in Agarose Plugs

Double stranded DNA breaks were detected as previously described. Briefly, cells were harvested by trypsinization, washed with PBS, and counted. 500,000 cells were embedded in 0.75% Clean Cut Agarose and allowed to solidify, before digesting overnight with Proteinase K at 50°C. The plugs were washed four times for 1 hour, before loading into a 1% agarose gel. The gel was run with 0.5X TBE at 14°C with a two-block program; block 1: 12 hour, 0.1 s initial, 30 s final, at 6V/cm; block 2: 12 hour 0.1 s initial, 5 s final, 3.8V/cm. The gel was then dried 2 hours at 50°C before staining with SYBR Green and imaging on a Typhoon.

### XRCC1 Recruitment and Analysis

RPE FAP-TRF1 cells were plated on coverslips so they would be ∼70% confluent the next day. They were then transfected with pEYFP-XRCC1 (1µg) and 6µl Fugene 6 (Promega) in Optimem (Gibco) using media without antibiotics. 24 hours later, the cells were treated with dye and light for 10 minutes, and then immediately subjected to CSK extraction before fixation. After washing, cells were mounted without DAPI. Only YFP positive cells were imaged and the CFP channel was used to mark telomeres (FAP-mCER-TRF1).

### Image Acquisition and Analysis

All IF images were acquired on a Nikon Ti inverted fluorescent microscope equipped with an Orca Fusion cMOS camera, or CoolSNAP HQ2 CCD. Z-stacks of 0.2 μm were acquired for each image and deconvolved using blind, iterative methods with NIS Elements AR software. For co-localizations, deconvolved images were converted to Max-IPs and converted to a new document. The object counts feature in NIS AR was used to set a threshold for foci that was kept throughout the experiment. The binary function was used to determine the intersections of 2 or 3 channels in defined regions of interest (ROI) (DAPI stained nuclei). For whole nuclei signal intensity, the automated measurements function was used on ROIs.

### Western Blotting

Cells were collected from plates with trypsin, washed, and then lysed on ice with RIPA buffer (Santa Cruz) supplemented with PMSF (1nM), 1x Roche Protease and Phosphatase Inhibitors, and Benzonase (Sigma E8263; 1:500) for 15 minutes and then incubated at 37°C for 10 minutes, before spinning down at 13,000 rpm for 15 minutes at 4°C. Protein concentrations were determined with the BCA assay (Pierce) and 10-30 μg of protein was electrophoresed on 4-12% (or 12% for OGG1ko blot) Bis-Tris gels (Thermo) before transferring to PDVF membranes (GE Healthcare). Membranes were blocked in 5% milk and blotted with primary and secondary HRP antibodies. Signal was detected by ECL detection and X-ray film.

### Reverse Transcription qPCR

RNA was extracted from cells using the Qiagen RNeasy Plus Mini kit. 500-1000 ng RNA was converted to cDNA using the High capacity RNA-to-cDNA Kit (Thermo). 50 ng cDNA was subjected to real-time qPCR using Taqman probes at 1x and the Taqman Universal PCR kit (Thermo). Data were analyzed using the delta delta Ct method.

### Flow Cytometry

To analyze apoptosis, cells were treated as indicated, and allowed to recover for 4 days. Floating cells were collected, and then attached cells were collected with trypsin and combined. After spinning down and washing, the cells were incubated with Alexa Flour 488 annexin V and 1 µg/ml propidium iodine in 1x annexin-binding buffer for 15 minutes in the dark (Thermo). After resuspending in additional binding buffer, the cells were analyzed on an Accuri C6 (Beckman) using FL1 and FL3.

For cell cycle analysis, 23 hours after treatment, cells were pulsed with 20 µM EdU, and incubated for an additional hour (Thermo). Cells were collected with trypsin, washed with 1% BSA in PBS, and then fixed with Click-IT fixative D. After washing with 1% BSA in PBS, the cells were permeabilized with 1x component E for 15 minutes, before performing Click chemistry with Alexa Flour 488 azide for 30 minutes in the dark. Cells were washed with 1x component E, and then resuspended in 500 µl FxCycle PI/RNase (Thermo) for 15 minutes before analyzing on Accuri C6.

### Seahorse Analysis

OCR and ECAR were measured using a SeahorseXF96 Extracellular Flux Analyzer (Seahorse Bioscience) essentially as previously described (Qian et al., 2019). After treatment and recovery for the indicated times, cells were seeded in XF96 cell culture plates at 8 × 10^4^ cells per well in the presence of Cell-Tak cell and tissue adhesive. Cells were then washed, and growth medium was replaced with bicarbonate-free medium. Thereafter, cells were incubated for another 60 min in a 37 °C incubator without CO_2_ followed by simultaneous OCR and ECAR measurements.

### Bulk RNA-Seq

RNA was prepared using the Qiagen RNAeasy Mini Plus kit. 1 μg total RNA was sent to Genewiz for library preparation and sequencing. RNA with a RIN >9 was polyA selected and fragmented before cDNA synthesis. Adaptors were ligated, PCR enriched, and then sequenced on a HiSeq 2x150 in paired end mode. Each sample was sequenced to at least 30 million reads.

The triplicate measurements of gene expression in the mRNAseq data was quantified using Salmon (0.7.2) to the HG19 refseq transcript annotations (Patro et al., 2017). Unique genes were obtained by summing across transcript isoforms and gene count matrixes from untreated and treated (‘dye + light’) and were analyzed with DEseq2 to obtain fold change and p-value scores for each gene (Love et al., 2014). Differentially expressed genes were defined as >0.5 log2 FC, -log10(pval)>10^4, and quantifiable (“expressed”, >10 counts) in all three cell lines. To determine if gene expression was altered in chromosomal regions nearby the telomeres (which may be exposed to oxidative stress from the FAP system) we binned genes by the distance of their start site from the chromosomal ends and averaged across genes of a given distance from the chromosome end. Gene set enrichment was preformed using FGSEA and Hallmark gene sets (Liberzon et al., 2015).

### Telomere Restriction Fragment (TRF) Analysis

TRF analysis was performed as previously described (Fouquerel et al., 2019). Briefly, genomic DNA was extracted from cells using Qiagen Tip-20 or 100 according to manufacturer’s instructions. The DNA was digested with HinfIII and RsaI overnight, before pulsed-field gel electrophoresis. After drying the gel, the molecular weight ladder was detected with Sybr Green (Thermo) and then hybridization with ^32^P labeled telomere probe was carried out as described.

### Telomere Shortest Length Assay (TeSLA)

Assay was performed as previously described with some modifications (Lai et al., 2017). Genomic DNA (50 ng) was ligated to TeSLA-T oligo before digestion with CviAII to create 5’ AT overhangs. DNA was further digested wit BfaI, NdeI, and MseI to generate 5’ TA overhangs (NEB). After dephosphorylation, AT and TA adaptors were ligated, and then PCR (4 per sample) was performed with AP and TeSLA-TP primers. PCR product was cleaned up using Genejet PCR Purification kit before electrophoresis. Telomere fragments were detected after drying the gel using in-gel hybridization as previously described (Fouquerel et al., 2019).

### Metaphase IF (Meta-TIFF)

Cells were collected by trypsinization, washed with PBS, counted and spun down. 200,000 cells were swelled in 0.2% potassium chloride and sodium citrate for 5 min at 37 °C and then cytocentrifuged onto slides (10 min, 2000 rpm, medium acceleration). Cells were fixed 4% formaldehyde in PBS, and then processed for IF/FISH as described above.

### Detection of Mitotic DNA Synthesis

16 hours after treatment, cells were incubated with 7 μM Cdk1 inhibitor RO3306 (Millipore) for 24 hours. Cells were washed with PBS, and then released into media with 20 µM EdU and colcemid for 1 hours before harvesting by mitotic shake-off. Metaphase spreads were prepared as described above and EdU staining performed using Click-iT™ EdU Alexa Fluor™ 594 imaging kit (ThermoFisher) after FISH staining.

