## Supplementary figures and images for "Telomeric 8-Oxoguanine Drives Rapid Premature Senescence In The Absence Of Telomere Shortening"

### Figure S1

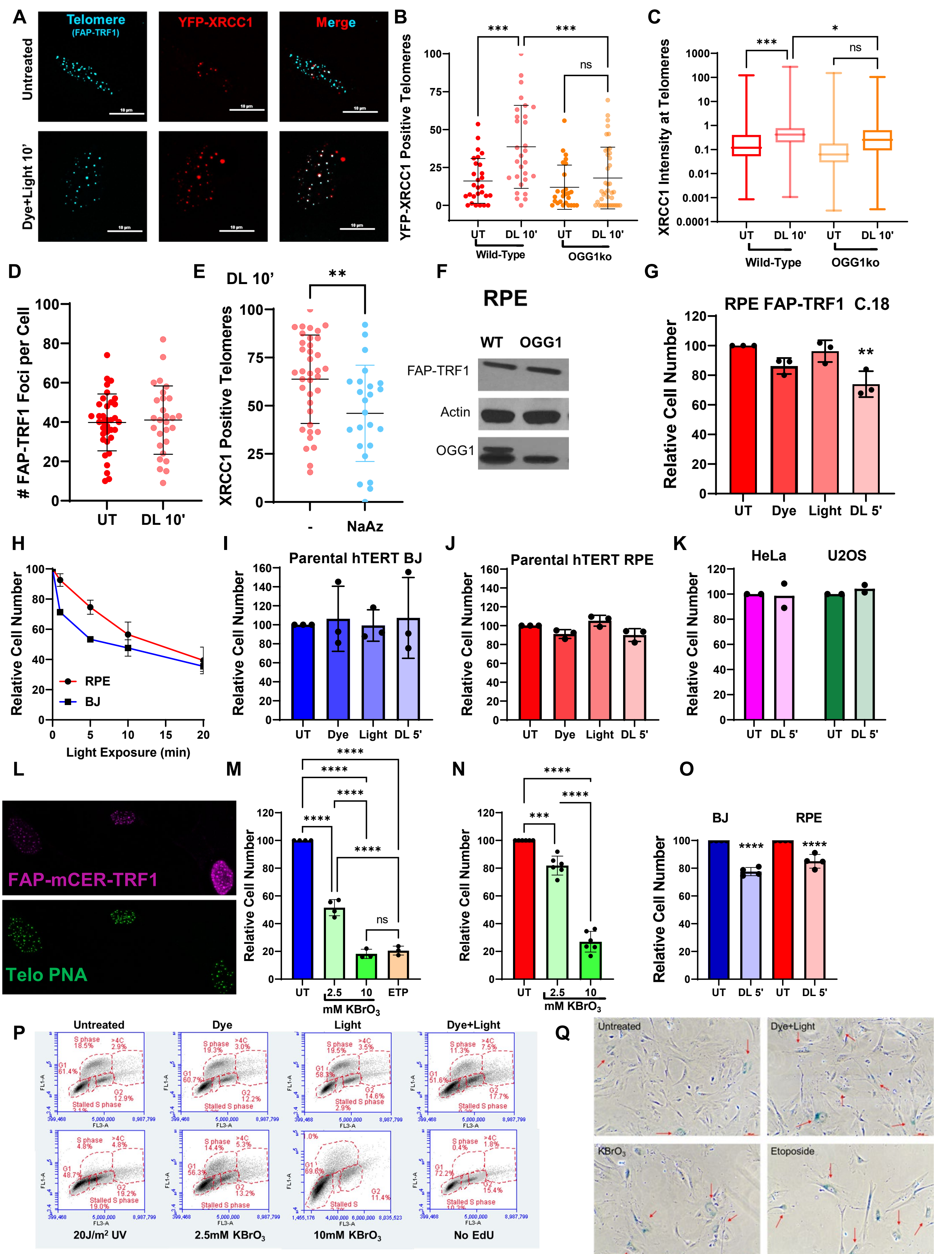

Supplemental Figure 1

### Figure S2

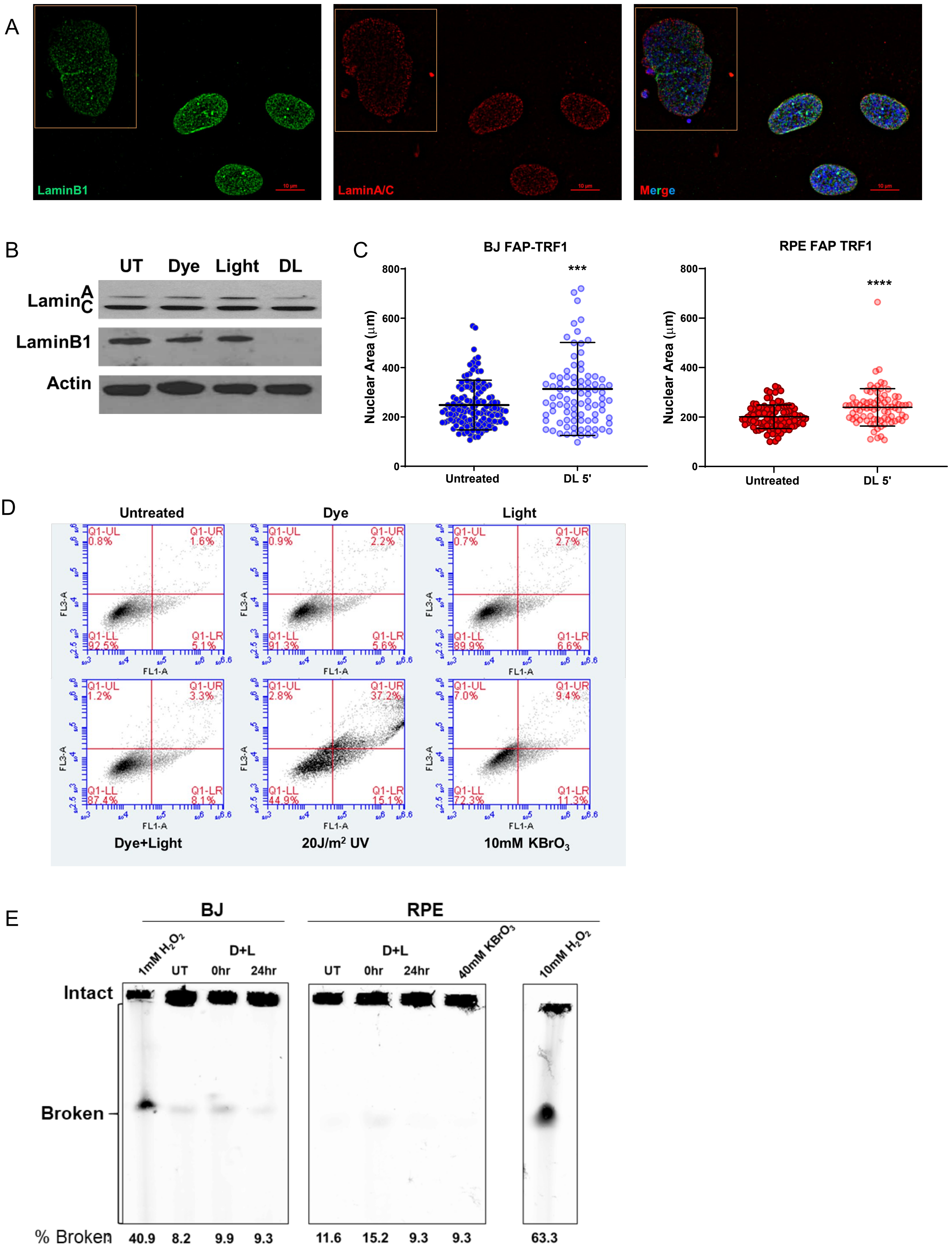

**Supplemental Figure 2**

### Figure S3

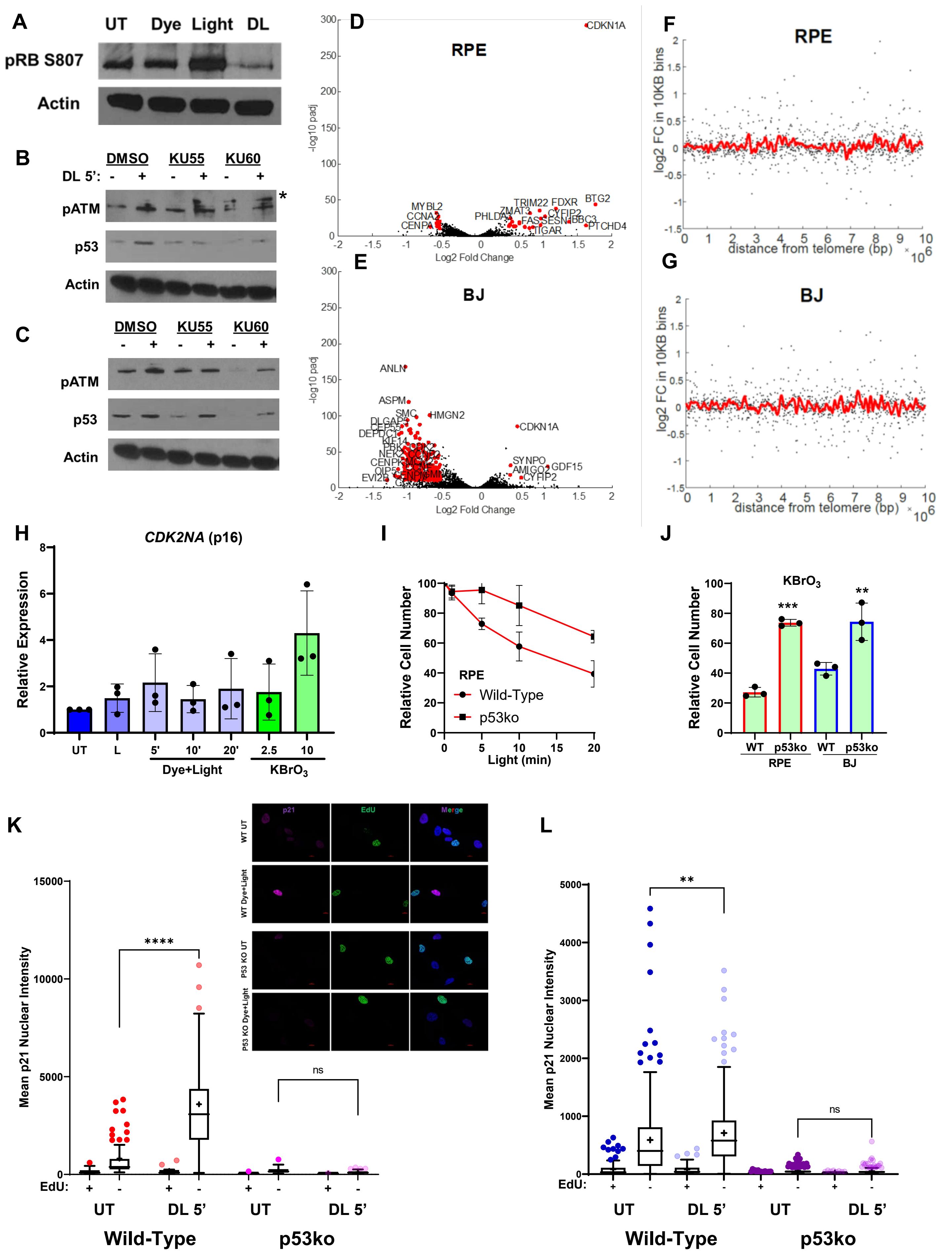

Supplemental Figure 3

### Figure S4

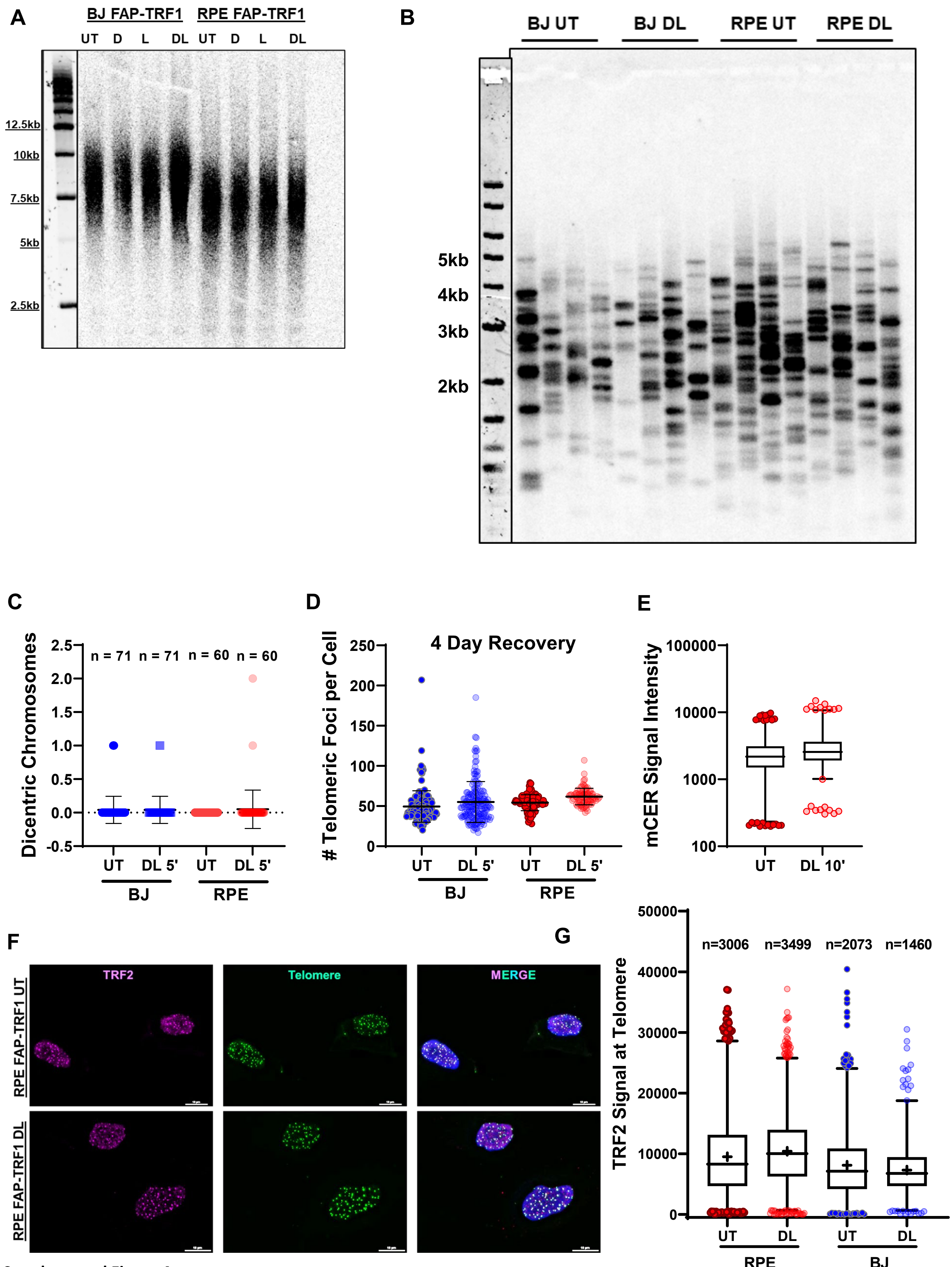

Supplemental Figure 4

### Figure S5

A

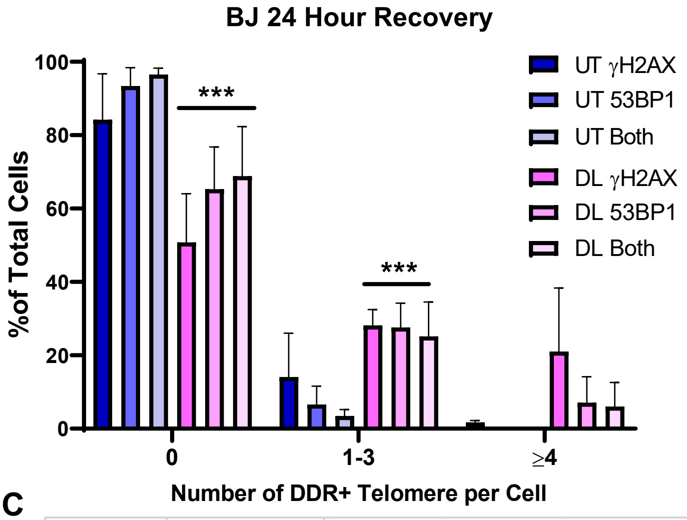

B

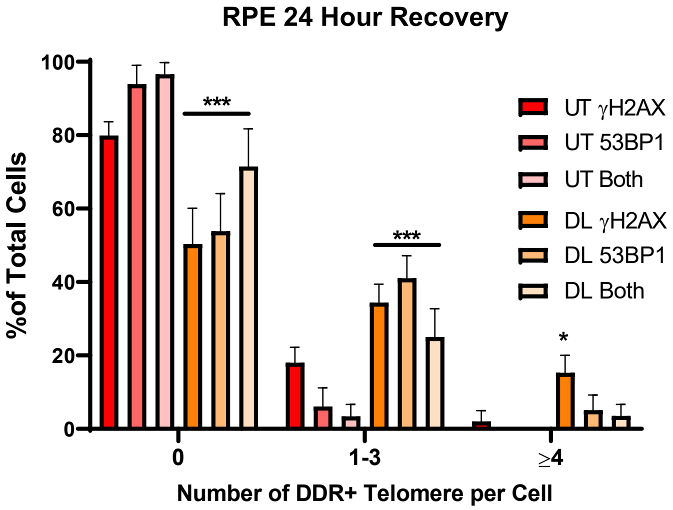

C

| | | Untreated | | | | DL 5' | | |
|---|---|---|---|---|---|---|---|---|
| | Experiment | γH2AX | 53BP1 | Sum | | γH2AX | 53BP1 | Sum |
| BJ | 1 | 1.43 | 0.00 | 1.43 | | 40.68 | 15.25 | 55.93 |
| | 2 | 2.33 | 0.00 | 2.33 | | 14.29 | 2.86 | 17.14 |
| | 3 | 1.43 | 0.00 | 1.43 | | 8.06 | 3.23 | 11.29 |
| | | | | 1.73 | | | | 28.12 |
| RPE | 1 | 0.79 | 0.00 | 0.79 | | 20.00 | 9.41 | 29.41 |
| | 2 | 5.38 | 0.00 | 5.38 | | 15.29 | 4.71 | 20.00 |
| | 3 | 0.00 | 0.00 | 0.00 | | 10.47 | 1.16 | 11.63 |
| | | | | 2.05 | | | | 20.35 |

D

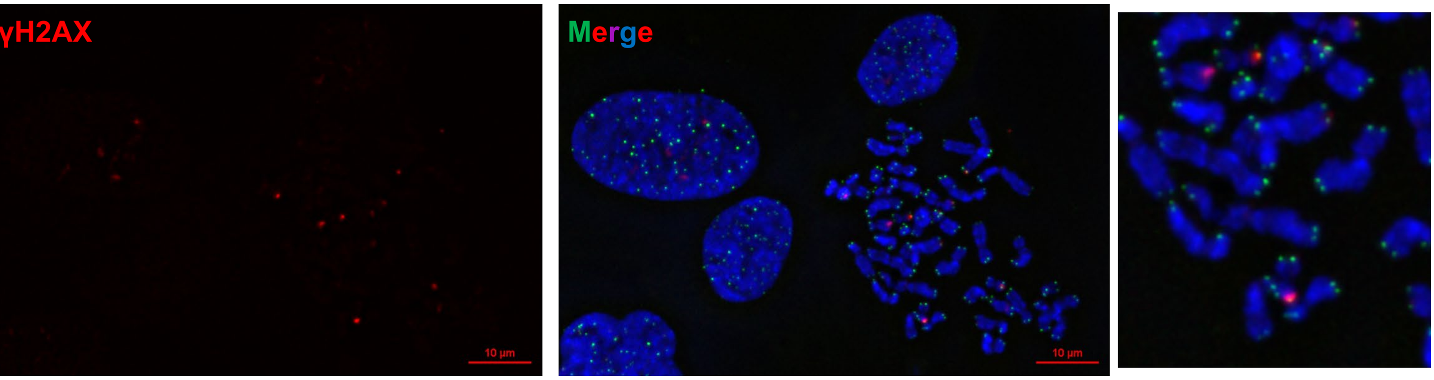

E

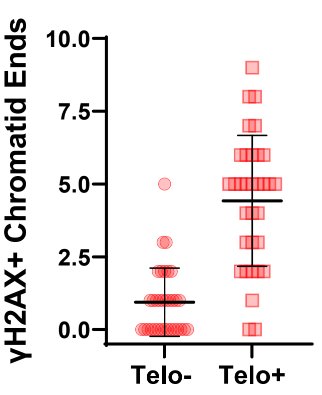

F

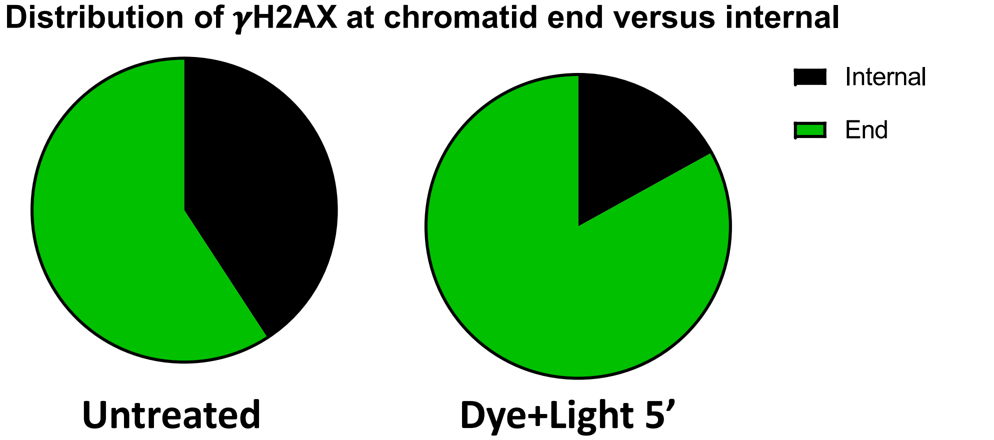

Supplemental Figure 5

### Figure S6

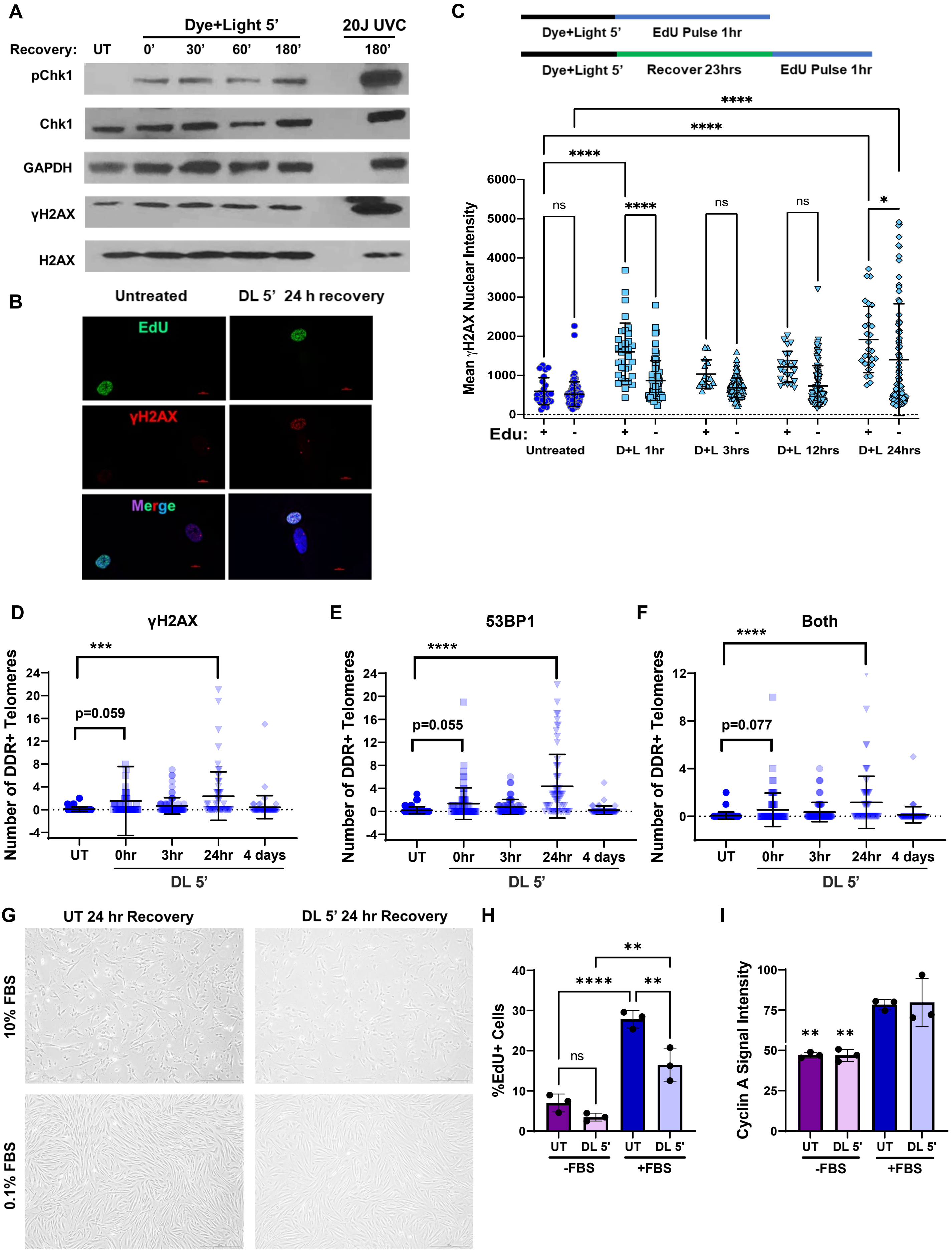

Supplemental Figure 6

### Figure S7

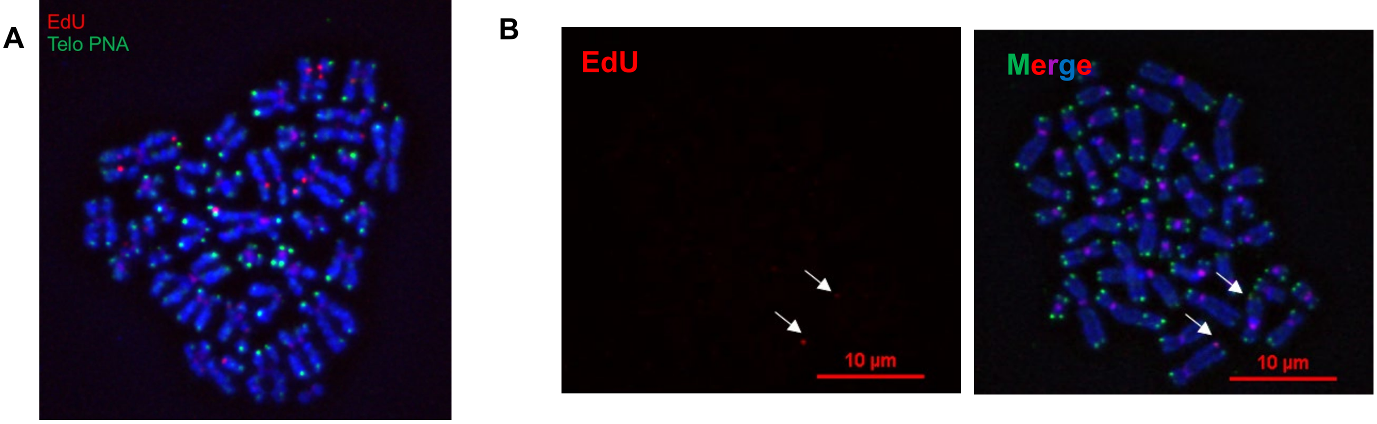

**Supplemental Figure 7**
