## Supplemental Table 1 for "Telomeric 8-Oxoguanine Drives Rapid Premature Senescence In The Absence Of Telomere Shortening"

| Hallmark | RPE | | BJ | |
|---|---|---|---|---|
| | padj | NES | padj | NES |
| E2F Targets | 2E-42 | -3.161 | 9.1E-58 | -3.0281 |
| $G_2$/M Checkpoint | 4.9E-38 | -3.0811 | 8.9E-50 | -2.945 |
| Mitotic Spindle | 9.3E-13 | -2.3133 | 2.4E-17 | -2.3424 |
| MYC Targets V1 | 0.00481 | -1.5783 | 5.5E-13 | -2.2066 |
| MYC Targets V2 | - | - | 0.00621 | -1.7299 |
| DNA Repair | - | - | 0.00027 | -1.74 |
| Spermatogenesis | 0.00045 | -1.9472 | 1.4E-05 | -2.0434 |
| Estrogen Response Late | 0.00147 | -1.7133 | - | - |
| p53 Pathway | 6.6E-19 | 2.8275 | 4.4E-05 | 1.80522 |
| Apoptosis | 2.5E-05 | 2.07155 | - | - |
| TNFα Signaling Via NFκB | 0.00038 | 1.85376 | - | - |
| Oxidative Phosphorylation | 0.00208 | 1.69939 | - | - |
| Interferon Gamma Response | 0.0022 | 1.73692 | - | - |
| Inflamatory Response | 0.00499 | 1.73654 | - | - |

**Supplemental Table 1**
